# Stochastic failure accumulation as a foundation for exponential mortality and selective disappearance

**DOI:** 10.64898/2026.05.25.727614

**Authors:** Ananda Shikhara Bhat, Hanna Kokko

**Affiliations:** Institute of Organismic and Molecular Evolution (iomE), Johannes Gutenberg University, 55128 Mainz, Germany; Institute for Quantitative and Computational Biosciences (IQCB), Johannes Gutenberg University, 55128 Mainz, Germany

**Keywords:** Gompertz-Makeham, senescence, reliability theory, frailty, damage accumulation, biological age

## Abstract

Most organisms become increasingly likely to die as they grow older, a phenomenon known as demographic senescence. We develop a general mathematical framework for modelling senescence in which individuals are viewed as comprising a fixed number of interacting intra-organismal sub-systems, each characterised by stochastic failure and repair rates. In many organisms, ‘failure begets failure’ because sub-systems are typically interdependent. We use diffusion approximations to derive an expression for population-level mortality curves in this setting, and demonstrate that greater interdependence leads to faster rates of senescence. Since individuals who die can no longer age, observed cohorts become increasingly composed of ‘lucky’ individuals that avoided death by (stochastically) taking paths in failure space associated with lower mortality despite no intrinsic differences in their quality. Selective disappearance of ‘unlucky’ individuals generates deviations of population-level measurements from individual-level trajectories, causing late-life mortality plateaus. We show that while these deviations must always exist, they may often be too small to detect, either because the failure accumulation process is stereotyped or because detection requires unreasonably large cohort sizes. We provide necessary and asymptotically sufficient conditions for Gompertz-Makeham mortality in our framework, with falsifiable predictions in terms of a scaling law for mortality near complete failure. We also show that the simplest null model in our framework leads to exact Gompertz-Makeham curves. Our work establishes a general framework for modelling demographic senescence and selective disappearance from biological first principles, and provides concrete predictions for when Gompertz-Makeham mortality emerges in such models.

## Introduction

Senescence, or ageing, is defined as a persistent increase in mortality or decrease in fecundity from the time since the birth of an organism (Shefferson et al., 2017). Explaining the prevalence of senescence across the tree of life is a classic problem in evolutionary biology (Medawar, 1952). Classic evolutionary theories of senescence crucially rely on either age-specific mutations to fecundity and survival (Williams, 1957; Hamilton, 1966) or strong trade-offs between the maintenance of somatic function and germline integrity (Kirkwood, 1977). However, they do not address the mechanistic, physiological basis for observed senescence patterns in any particular organism, and typically make neither lifespan predictions nor derive expectations for mortality curves (Lehtonen, 2020).

An alternative, mechanistic approach to understanding senescence considers the accumulation of damage, or ‘failures’, of physiological processes required for organismal function (Gavrilov and Gavrilova, 1991; Bega and Hadany, 2026). In this view, an organism comprises many interdependent biological processes, each with some intrinsic risk of failure, and senescence arises via a catastrophic propagation of failures between interdependent subsystems (Gavrilov and Gavrilova, 1991; Boonekamp et al., 2015). As is typical in complex systems theory, there is considerable flexibility in the choice of what constitutes a subsystem (Ledberg, 2020; Nielsen et al., 2024): intra-organismal sub-systems could be genes, molecular interaction pathways, tissues, or physiological systems. The crucial requirements are that there are many of them, that they interact with each other, and that they are important for survival and mortality. Farrell et al. (2021), Cohen et al. (2022), and Meyer et al. (2025) provide conceptual overviews of the complex systems approach to ageing; also see López-Otín et al. (2023) for a further list of biologically motivated candidates for sub-systems.

Classic reliability theoretic models of senescence arrange intra-organismal subsystems either ‘in series’, ‘in parallel’, or as a combination of parallel blocks connected in series (Gavrilov and Gavrilova, 2001; Laird and Sherratt, 2009, 2010; Boonekamp et al., 2015). More complicated dependency structures in terms of networks have also been proposed (Rosen, 1958; Vural et al., 2014; Sun et al., 2020; Nielsen et al., 2024), including data-driven computational models of failure/-damage accumulation (Farrell et al., 2021). Contrasting with this extensive effort, the mapping from failures to mortality has received relatively scant attention. Stochastic inter-individual heterogeneity in the damage accumulation process is similarly absent in reliability-theoretic models of senescence, despite recent work showing its importance for predicting population-level patterns of demographic senescence (Meyer and Schumacher, 2024) and life-history theory (Snyder et al., 2021). Most mortality models incorporating inter-individual heterogeneity instead do so by assigning individuals to fixed ‘frailty’ classes that differ in ‘quality’, typically affecting ageing rate (Vaupel et al., 1979; Vaupel and Missov, 2014; Balan and Putter, 2020). However, stochastic variation accumulates with age (Meyer and Schumacher, 2024) and can thus cause dynamically varying heterogeneity between individuals (Tuljapurkar et al., 2009; Snyder et al., 2021). A general framework in which population-level mortality patterns are systematically derived from the processes of physiological ageing due to stochastic sub-system failure (damage accumulation) at the individual level is thus desirable (Farrell et al., 2021; Cohen et al., 2022).

We introduce a flexible modelling framework which derives both stochastic failure/damage accumulation dynamics at the organismal level, and observed mortality patterns at the population level, from intra-organismal first principles. We first derive results for a simple model where failure accumulation follows a stochastic birth-death process and individuals are assigned mortalities that are a function of accumulating failures, but we explicitly exclude selective disappearance as the mortalities do not translate into actual deaths. We call this initial model the ‘Tithonus model’ after the myth of Tithonus from Greek mythology (Graves, 2017, chapter 40). We then develop a second, more complete model in which mortality is implemented as a true stochastic rate and failure accumulation patterns are thus the result of a birth-death process with killing (Steinsaltz and Evans, 2006), but which can be analysed using the simpler Tithonus model.

Using these models and their diffusion approximations, we present general analytical expressions for the expected population-level mortality curve. By representing measurements made in our complete model in terms of measurements made on the Tithonus model, we show that selective disappearance naturally and unavoidably emerges due to stochastic heterogeneity in the failure accumulation process and causes biases in population-level measurements. We demonstrate that such selective disappearance causes late-life mortality plateaus, and study when these plateaus are difficult to detect in empirical data.

We also postulate some criteria that biologically realistic functional forms of failure accumulation and mortality rate functions should obey, and find that a simple null model satisfying these postulates predicts exact Gompertz-Makeham mortality (in the Tithonus model). Going beyond the null model, we provide a necessary condition for Gompertz-Makeham mortality and a sufficient condition for asymptotically Gompertzian mortality in our framework. These necessary and sufficient conditions take the form of scaling laws for the dependence of mortality on failures near complete failure, and are thus concrete experimental predictions of our framework when the Gompertz-Makeham law is known to be a good description of population-level mortality patterns.

### The Tithonus model: Modelling failure accumulation and mortality separately

Tithonus in ancient Greek mythology was a mortal lover of Eos, Goddess of the Dawn. After much pleading from Eos, Zeus granted Tithonus immortality so he could be with Eos forever. However, Eos neglected to also request Tithonus remain healthy and youthful. Tithonus thus withered away, yet, no matter how feeble and decrepit, he could not die (Graves, 2017, chapter 40). We model Tithonus-style lives, intentionally ignoring the contradiction of deriving mortality rates while also assuming immortality. The intention is to achieve a clean separation of failure accumulation and resultant mortality risk, as examining them separately allows us to understand when to expect Gompertz-Makeham curves.

We consider a population of independent, identical, non-interacting individuals (all called Tithonus, to make it clear throughout the text when we are operating within this assumption system). Tithonus consists of *N* intra-organismal sub-systems, is pristine at birth (*F*_0_ ≥ 0 failed sub-systems), and sub-systems stochastically fail thereafter. We assume that failure begets failure: in a Tithonus with *F* failed sub-systems, additional sub-system fails occur at rate *R*_+_(*F*). A failed sub-system is independently repaired at rate *R*_−_(*F*). We assume *R*_+_(*F*) − *R*_−_(*F*) > 0 for every 0 ≤ *F* < *N*, i.e., the number of failed sub-systems is always more likely to increase than decrease (see Discussion for relaxing this assumption). This reflects the view that wear-and-tear and imperfect repair (Gorbunova et al., 2007) together make it difficult to keep organisms pristine. We further assume that the failure rates depend on the *proportion* of failed sub-systems, and that the total number *N* only rescales this dependence. This is achieved by assuming *R*_±_(*F*) = *Nr*_±_(*F*/*N*), where *r*_±_ are *O*(1) functions.

We will thus track the proportion *f* := *F*/*N* rather than the total number of failed sub-systems *F*. For conciseness, we use the term ‘failures’ for the proportion *f*, and note that it obeys the transitions

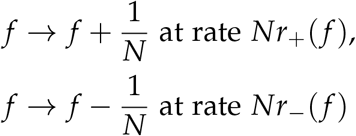

We call the expected net rate of failure accumulation in a Tithonus with *f* failures (*r*( *f* ) := *r*_+_( *f* ) − *r*_−_( *f* ) > 0) the *failure accumulation rule* of the model. The quantity *τ*( *f* ) := *r*_+_( *f* ) + *r*_−_( *f* ) quantifies the total rate at which an individual with *f* failures experiences a change in failures. Following work in other areas of population biology, we will call *τ* the *turnover rate* (Bhat, 2025; Bhat and Guttal, 2025; Komarova and Wodarz, 2025). See Appendix A for a table of notation used in the main text.

#### Failure dynamics in the Tithonus model

Let f_*t*_ denote the stochastic process describing the proportion of failures accrued by a Tithonus of age *t*. We use a system size expansion (‘diffusion approximation’ in population biology) to show (Supplementary section S1) that if *N* is not too small, f_*t*_ is approximated by the solution to the Itô stochastic differential equation (SDE)

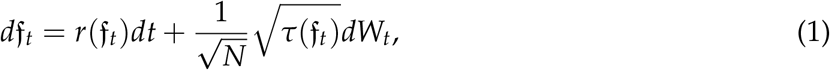

where *W*_*t*_ is a Wiener process (standard Brownian motion).

We now define the ‘mortality rule’ of the model, *µ*, such that *µ*(*f*) describes the mortality assigned to Tithonus with *f* failures (without actually causing death). We now use Itô’s formula (Itô, 1951) to find the mortality rate of Tithonus at age *t* obeys the stochastic differential equation

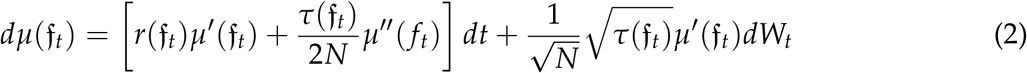

Here, *µ*^′^(*f*) and *µ*^′′^(*f*) are the first two derivatives of the mortality rule evaluated at the current value of *f* ; they respectively indicate whether mortality increases or decreases with *f*, and whether this dependence is (locally) accelerating or decelerating.

As *N* → ∞, Eq. 2 becomes the ordinary differential equation 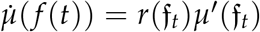, and only the first derivative *µ*^′^ of the mortality rule remains important. When *N* < ∞, the second derivative also plays a role (intuitively due to Jensen’s inequality, see e.g. Box 3 in Bhat and Guttal (2025)). For instance, in an accelerating mortality rule (*µ*^′′^(*f*) > 0), the expected mortality curve E[*µ*(*f*_*t*_)] predicted by Eq. 2 is always higher than the curve predicted in the *N* → ∞ limit (Fig 2C,D), an effect that becomes weaker As *N* increases.

### The complete model with selective disappearance

The simplicity of the Tithonus model comes at a cost of assigning mortality rates to individual states (failures) without taking into account that individuals who die can no longer age. In reality, the probability of reaching some states (failure values) alive may be low or zero. To better understand the shortcomings of the Tithonus simplification, we now present a more complete model that allows mortality to act at the appropriately stochastic rate for each individual (no longer called Tithonus). This allows us to study how the mortality process itself feeds back, via the disappearance of individuals, into predicting the mortality curve expected to be observed at the population level.

We model the failure dynamics 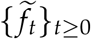 as a birth-death process with killing (Karlin and Taylor, 1981, p. 161) on 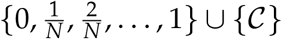 with transition rates

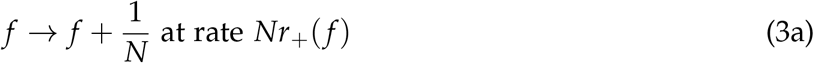

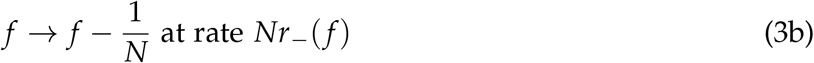

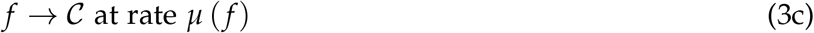

Here, *C* is a ‘cemetery state’ (a term borrowed from human demography) used to denote the death of the individual (Karlin and Taylor, 1981, p. 161). The Markov process (Eq. 3) can be simulated exactly using the Gillespie algorithm (Gillespie, 1977; Kokko, 2024).

If *N* is not too small, a diffusion approximation (supplementary S1) can be used to approximate the exact discrete state process defined by Eq. 3 by a continuous state stochastic process called a *killed diffusion* (Karlin and Tavaré, 1982; Steinsaltz and Evans, 2006). The failure accumulation can be characterized as the probability distribution of observed failures at age *t*, conditional on the individual having reached age *t* alive. In other words, if the random variable *T*_mort_ denotes the age of death, we wish to describe 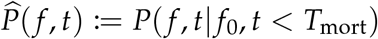, the probability that a living age *t* individual has a proportion *f* of failed subsystems given that at age 0 this proportion was *f*_0_. Yashin et al., 1985 (their Eq. 7 and their Appendix A; see our supplementary section S1) have proved that *P* ( *f, t*) obeys the partial differential equation

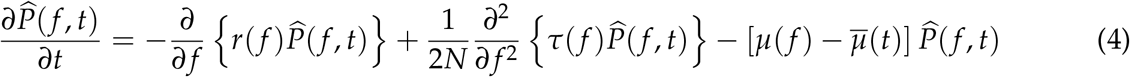

where

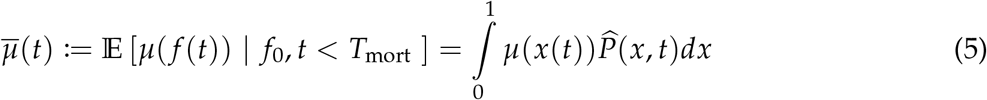

is the expected force of mortality conditioned on remaining alive.

While Eq. 4 describes how the probability distribution of failures as a whole changes over age, an alternative, ‘pathwise’ description lets us follow an individual through the course of its life and ask how its state (failures) and mortality risk are expected to change in terms of means and variances (see Fig 1 for a visual representation of the two different descriptions). We provide such a weighted paths model below.

**Figure 1:**
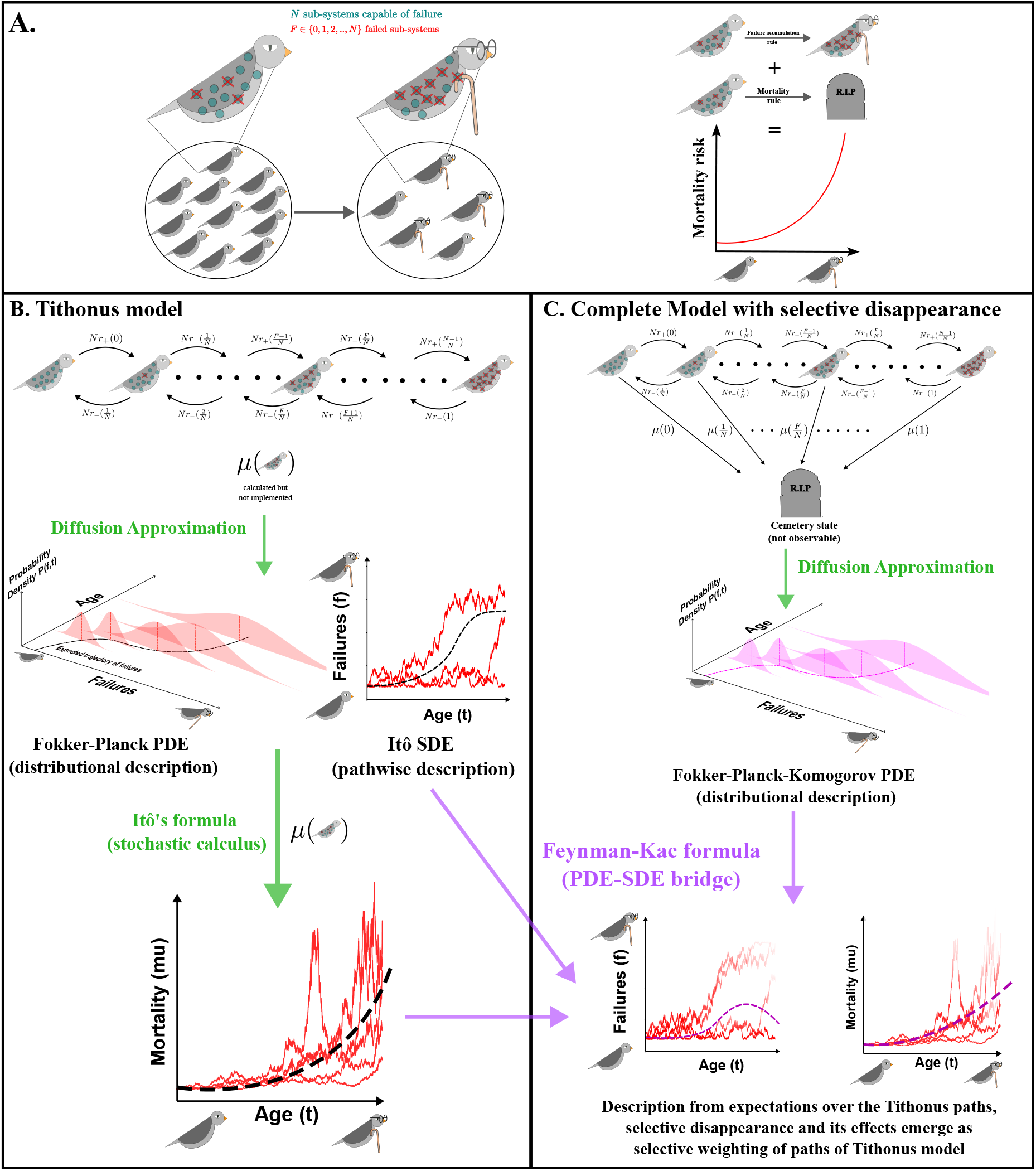
An outline of the general approach of this paper. **(A)** We model a large population of independent, non-interacting organisms. Senescence at the individual level occurs as intraorganismal sub-systems fail and impact each other’s failure rate. Our goal is to understand how failure accumulation and mortality together predict mortality curves, which in this framework are an emergent pattern. **(B)** The Tithonus model focuses on the failure dynamics alone, allowing to investigate relationships between age, failures and mortality. An individual in which 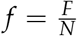 sub-systems have failed has a failure rate *Nr*_+_(*f*) and a repair rate *Nr*_−_(*f*). We calculate mortality according to some mapping between failures and mortality, but no individuals actually die. Here, a direct ‘pathwise’ description of that follows focal individuals through the course of their life is possible. **(C)** In our complete model, mortality is incorporated along with failure and repair as a stochastic transition rate at the individual level; the original state space is augmented with an additional, unobserved ‘cemetery’ state for dead organisms. An individual in which 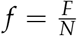 sub-systems have failed has a failure rate *Nr*_+_(*f*), a repair rate *Nr*_−_(*f*), and a mortality rate *µ*(*f*). For the complete model, the pathwise description is in terms of paths of the Tithonus model.

**Figure 2:**
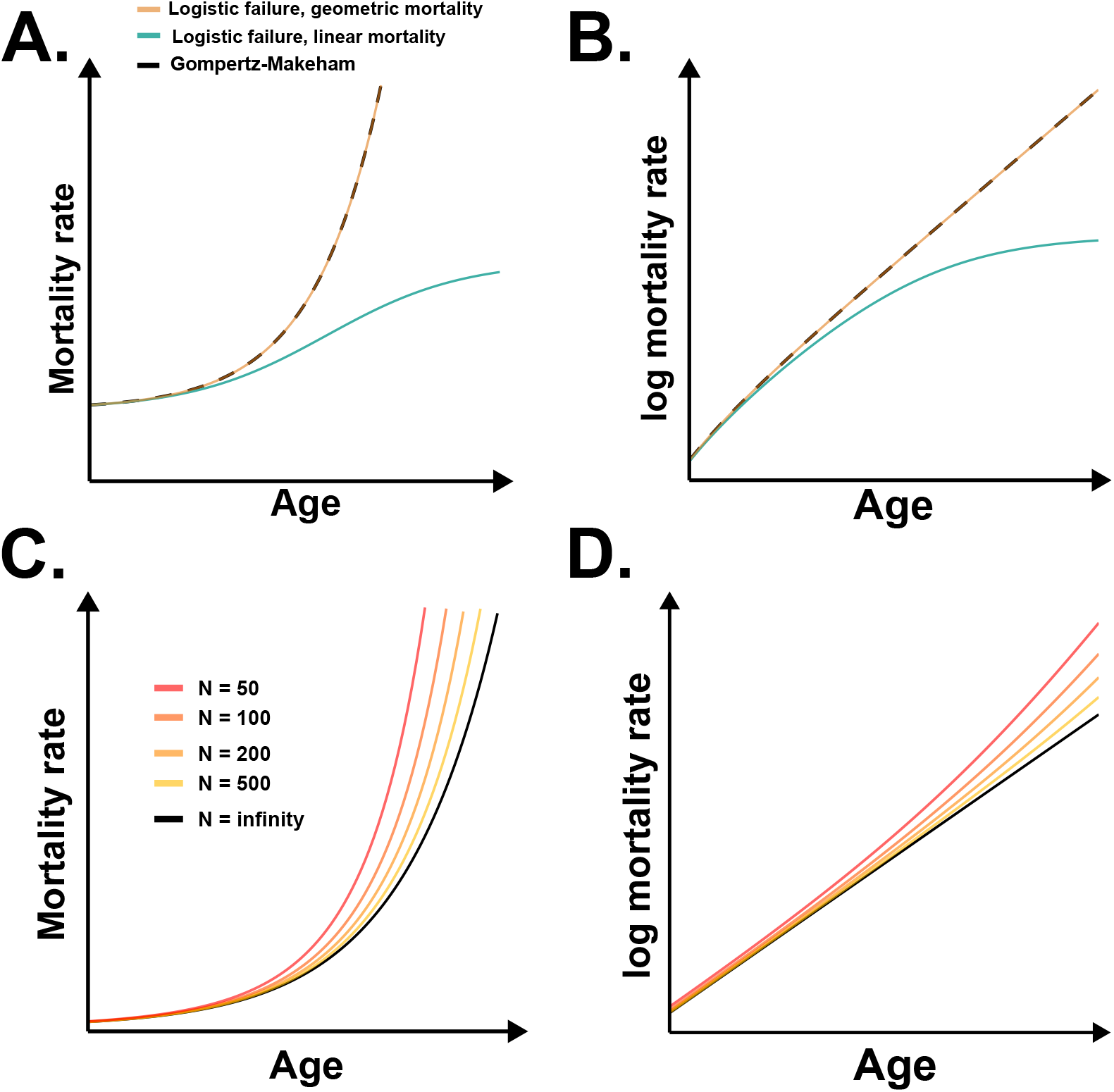
The predictions from the Tithonus model with logistic failure accumulation (Eq. 12). In all plots, we use *r*_+_( *f* ) = (1 − *f* )(2*ϕ*_*ρ*_ + *k*_*ρ*_ *f* ) and *r*_−_( *f* ) = (1 − *f* )*ϕ*_*ρ*_, so that the failure accumulation rule is *r*( *f* ) = (1 − *f* )(*ϕ*_*ρ*_ + *k*_*ρ*_ *f* ), *i.e*. the logistic rule Eq. 12. Panels **(A)** and **(B)** illustrate the *N* → ∞ predictions in comparison to a Gompertz-Makeham curve. If the mortality rule is linear (*µ*( *f* ) = *µ*_*e*_ + *χ f* ), the predicted mortality curve deviates from Gompertz-Makeham later in life, leading to a plateau. If we instead use the geometric mortality rule Eq. 13, the predicted mortality curve is exactly Gompertz-Makeham. Panels **(C)** and **(D)** illustrate the expected behaviour of the Tithonus model with the geometric mortality rule for finite *N*, from the expectation of Eq. 2. The mortality curve for finite *N* is uniformly higher than the *N* → ∞ curve. Axis ticks are deliberately left blank to underscore that quantitative values of ages and mortality hazards are arbitrary, because we are focused on the shape rather than the scale of the predicted mortality curve. Parameter values: **(A**,**B)** *ϕ*_*ρ*_ = 10^−3^, *k*_*ρ*_ = 0.1, *µ*_*e*_ = 10^−5^, *χ* = 1, *f*_0_ = 0.001; **(C**,**D)** *ϕ*_*ρ*_ = 10^−3^, *k*_*ρ*_ = 0.1, *µ*_*e*_ = 10^−5^, *χ* = 0.2, *f*_0_ = 0.01.

### The Feynman-Kac formula reveals the emergence of selective disappearance

Suppose an individual, born with *f*_0_ failures at birth, accumulates failures and experiences mortality risk according to our complete model with selective disappearance (Eq. 4). We denote this stochastic process by 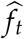. Let *M* : [0, 1] → R be a measurement made on an individual that bears *f* failures. For instance, choosing *M*(*f* ) := *f* measures failures, and choosing *M*( *f* ) := *µ*(*f*) measures the mortality associated with having *f* failures subject to the mortality rule *µ*. Other uses, such as *M* referring to reproductive output or nesting ability, also work, as long as the age-dependency of *M* is fully captured by failures *f* with no need to know chronological age per se (the technical term is that *M* is ‘time homogeneous’).

We are interested in 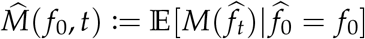, the value of the measurement performed on an individual that follows our complete model Eq. 4 and was born with 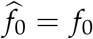 failures. Even though Eq. 4 is conditioned on the individual being alive, 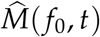 can be represented in terms of the paths (in failure space) of Tithonus, with paths in failure space that are more likely to be associated with death/mortality being systematically less likely to contribute to the measurement 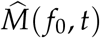. Intuitively, our result says that ‘unlucky’ individuals who have more failures than expected are more likely to die before reaching age *t* (see Snyder et al., 2021) and thus contribute less to the average value of the measurement at age *t*. More precisely, supplementary sections S2-S3 use a version (Øksendal, 1998, theorem 8.2.1(b)) of the Feynman-Kac formula (Feynman, 1948; Kac, 1949) to show that 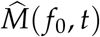 can be written as

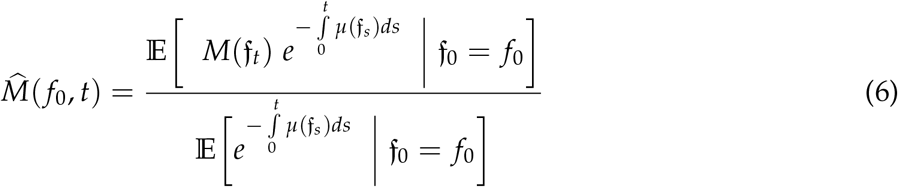

where f_*t*_ is the stochastic process describing the failures accrued by Tithonus at age *t* (Eq. 1) and the conditional expectation is over paths of {f_*t*_}_*t*≥0_ starting from *f*_0_ at *t* = 0 (see supplementary section S2.3 for an explanation of how to take conditional expectations over paths).

Since *µ*( *f* ) is the instantaneous mortality risk when an individual has *f* failures, 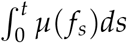 is the cumulative mortality risk associated with the entire path { *f*_*s*_}_*s*∈[0,*t*]_. The quantity 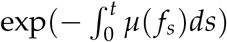, typically called survivorship, thus measures the probability of survival from age 0 to age *t*, when failure accumulation follows the path { *f*_*s*_}_*s*∈[0,*t*]_. The numerator of the RHS of Eq. 6 thus ‘sums up’ all possible values of the measurement *M* performed on Tithonus at age *t* (from Eq. 1), while weighting each possible measurement by the survivorship along the particular path the process took to arrive at its final value.

We show in supplementary section S3 that the denominator of Eq. 6 obeys

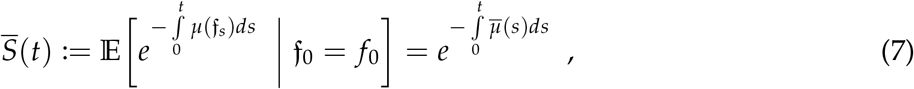

where 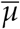 is given by Eq. 5. Intuitively, 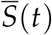 thus calculates the survivorship corresponding to the “average force of mortality” 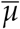 conditioned on individuals not dying. We also show in the supplementary (Eq. S30) that 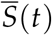 quantifies the expected number of individuals who reach age *t*. More precisely, if a population initially has *K* individuals of age 0, each individual following our complete model Eq. 3, then a proportion 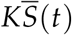 of them are expected to reach age *t* alive. Motivated by this, we henceforth call 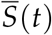 the “expected survivorship” in the population. Thus, the relation 6 clarifies the precise way in which selective disappearance affects measurements — those paths associated with death/mortality contribute systematically less to the value of measurement 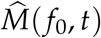.

Since the Tithonus model can be written as the solution to an SDE, it is both easier to handle analytically, and much more efficient to simulate, than the exact stochastic process Eq. 3. Our work therefore provides efficient numerical methods for deriving the effects of selective disappearance. The analytical Feynman-Kac predictions generally agree with direct individual-based Gillespie simulation of the exact stochastic process 3 (figure 3 provides an example). Both in turn differ significantly from Tithonus predictions that ignore disappearance.

**Figure 3:**
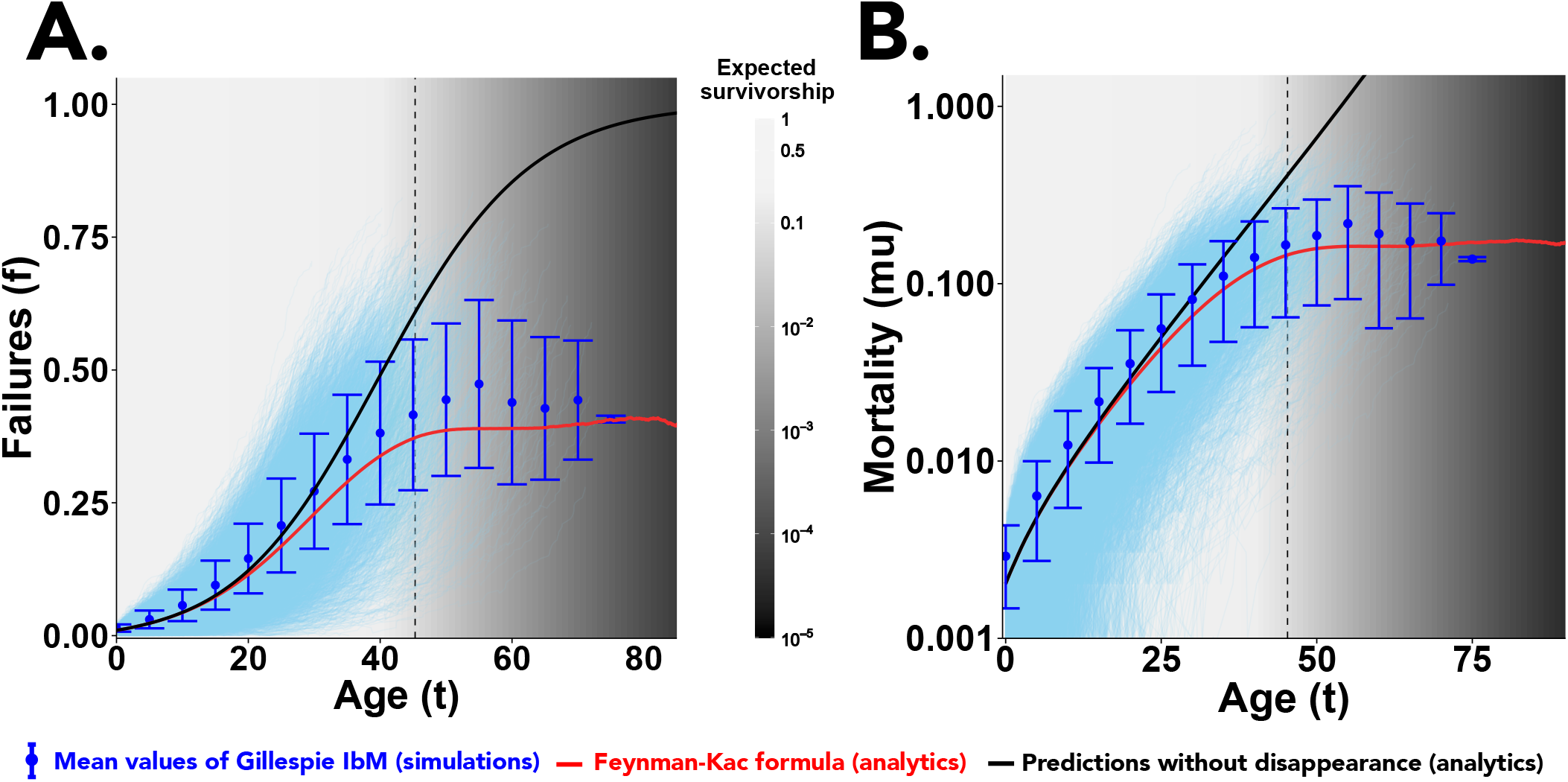
**(A)** Failure and **(B)** mortality accumulation curves predicted under the logistic failure accumulation rule Eq. 12 with the geometric mortality rule Eq. 13. Light blue trajectories are from 9500 Gillespie simulations of the exact stochastic process 3, and dark blue points (with error bars) are the mean values among these trajectories. Error bars are standard deviations. Black curve is the expected trajectory of the Tithonus model (Eq. 1 and Eq. 2), calculated by numerically simulating 20,000 realisations of the SDEs using the Euler-Maruyama method. Red curve is the Feynman-Kac formula (Eq. 6 with *M*( *f* ) = *f* for failures and *M*( *f* ) = *µ*( *f* ) for mortality) with expectations calculated using the numerical simulations of the Tithonus SDEs. Shade of the background represents expected survivorship from Eq. 7. Dotted vertical line indicates the point when 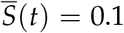. We use *r*_+_( *f* ) = (1 − *f* )(2*ϕ*_*ρ*_ + *k*_*ρ*_ *f* ) and *r*_−_( *f* ) = (1 − *f* )*ϕ*_*ρ*_, so that the failure accumulation rule is *r*( *f* ) = (1 − *f* )(*ϕ*_*ρ*_ + *k*_*ρ*_ *f* ) and the turnover is *τ*( *f* ) = (1 − *f* )(3*ϕ*_*ρ*_ + *k*_*ρ*_ *f* ). The means of the Gillespie simulations are generally higher than the Feynman-Kac prediction because rare, extremely ‘lucky’ individuals with few failures and exceptional lifespans are (by definition) unlikely to be present in any finite size sample of realisations of a stochastic simulation. Parameter values: *ϕ*_*ρ*_ = 10^−3^, *k*_*ρ*_ = 0.1, *µ*_*e*_ = 10^−5^, *χ* = 0.2, *N* = 200, *f*_0_ = 0.01.

The discrepancy between the Tithonus model and our complete model, which shows the effects of selective disappearance, becomes more pronounced at later ages. Importantly, however, survivorship also declines with age (background gradient in Fig 3), and this means that not many individuals necessarily survive up to the point where the deviation becomes noticeable. Fig 3 exemplifies this by illustrating the age (vertical dotted lines) at which only 10% of the initial cohort is alive. The deviation from Tithonus remains mild for the majority of individuals. Moreover, the survivorship plummets catastrophically past this point (notice that the gradient is on a logarithmic scale).

#### How much of a difference can selective disappearance make?

We have shown that selective disappearance can alter the observed values of measurements (Fig 3) from what would have been observed if there were no disappearance, but also that most individuals may be dead before the effect becomes stark. A natural follow-up question is whether ‘most are dead before deviations become clearly visible’ is a typical pattern. For instance, in humans, mortality beyond approximately 30 years of age fits the Gompertz curve well (see Figure 1 in Pichler and Uhlig, 2023 for a recent example in Germany), with no obvious deviation at oldest ages, and the existence of mortality plateaus in human demography is debated (see Discussion). Mathematically, the question concerns the expected difference between the measurement *M* made on Tithonus, and the corresponding measurement made on individuals following our complete model (the latter measurement being conditioned on the observed individual remaining alive), i.e. the value of 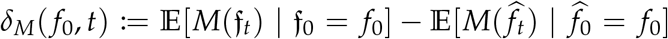 . Using Eq. 6 in the definition of *δ*_*M*_( *f*_0_, *t*) reveals that *δ*_*M*_ obeys

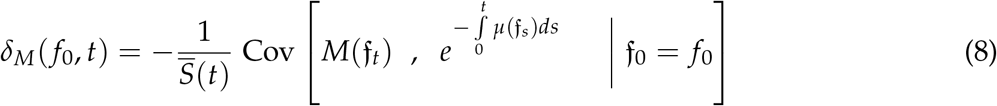

where Cov[*X*_*t*_, *Y*_*t*_|*X*_0_ = *x*_0_] := E[*X*_*t*_*Y*_*t*_|*X*_0_ = *x*_0_] − E[*X*_*t*_|*X*_0_ = *x*_0_]E[*Y*_*t*_|*X*_0_ = *x*_0_] is the conditional covariance process at time *t* between two stochastic processes {*X*_*t*_}_*t*≥0_ and {*Y*_*t*_}_*t*≥0_. Eq. 8 states that we expect selective disappearance to alter measurements significantly (|*δ*_*M*_| is large) if the measurement *M* is correlated with survivorship. If large values of *M* associate with low survivorship, the covariance is negative, and *δ*_*M*_ is positive: the expected value of the measurement is lower with than without disappearance. Conversely, if larger numerical values of *M* are associated with higher survivorship, the opposite statements apply. Furthermore, the magnitude of the difference, |*δ*_*M*_|, is proportional to the magnitude of covariance between the measurement and survivorship. In Supplementary section S4, we show that the absolute value of *δ*_*M*_ can be bounded above by

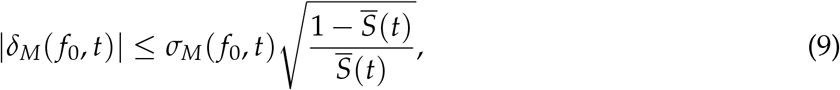

where 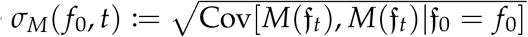 is the standard deviation process for the measurement *M*(f_t_), i.e. a quantification of the heterogeneity in the failure accumulation process {f_*t*_}_*t*≥0_ at age *t*. The term within the square root can be interpreted as an odds ratio: the denominator and the numerator are the expected survivorship and its complement, respectively.

Ineq. 9 can be used to show the effect of selective disappearance on mortality curves, if we assume that the turnover *τ*( *f* ) is bounded, *i.e*. that there is some finite number *τ*_max_ such that *τ*( *f* ) ≤ *τ*_max_ ∀ *f* ∈ [0, 1]. The boundedness is biologically justified: put informally, only so many things can happen within an organism per unit time. Let us further assume that *µ*^′^( *f* ), the rate of change of mortality as a function of failures, is also bounded. Using Eq. 2 corresponding to the putative mortality of Tithonus, supplementary section S4 shows that the effect of selective disappearance on mortalities, *δ*_*µ*_( *f*_0_, *t*), obeys

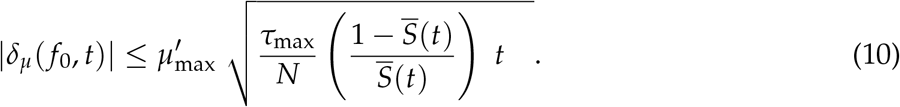

This shows that the effects of selective disappearance are limited by (a) 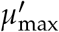, how quickly mortality can change as failures change (slower implies less visible effects of disappearance), (b) *t*, the age at which the measurement is made (the effects are less visible at smaller ages), (c) 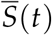, the overall expected survivorship at age *t* (greater survivorship implies less visible (but note that small cohort sizes at advanced ages also cause statistical sampling issues, see below) and (d) *N*, the total number of sub-systems that can fail. The effect of increasing *N* can be rephrased by noticing that a large *N* means that each individual is likely to follow a distinct ‘path to death’: the identity of the subsystem that fails first is then likely to differ between individuals, as does the identity of the ‘final straw’, and all else in between. We predict that the uniqueness of the causal route to each death weakens the signature of selective disappearance in any longevity dataset. An imaginary organism with *N* = ∞ would not show it at all, while an organism with only a small number of sub-systems capable of failure offers the best prospects for selective disappearance being observable in demographic datasets. Additional statements about the effects of factors such as extrinsic mortality can be made by choosing specific functional forms for *µ* (see supplementary section S5 for an example).

Eq. 8 shows that selective disappearance always reduces the population-level mortality rate — moving towards a plateau-like shape —-because the covariance between mortality and survivorship is negative. Further, the 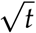 on the right hand side of Ineq. 10 indicates that *δ*_*µ*_ is smaller at younger ages, i.e. confirming the intuition that selective disappearance impacts mortality more strongly in later life (red curves and blue points in Fig 3). Theorem 3.4 in Steinsaltz and Evans (2006), together with Steinsaltz and Evans (2004), proves that at sufficiently advanced ages, the mortality curve arising from Eq. 4 (or, equivalently, Eq. 6 with *M*( *f* ) = *µ*( *f* )) will plateau in a strict sense, *i.e*. remain strictly constant with age.

Bounds such as Ineq. 10 concern the maximum value that *δ*_*µ*_ is mathematically allowed to take at age *t*. It can thus be thought of as an estimate of the ‘opportunity’ or ‘potential’ for selective disappearance to be important. Importantly, the bound is calculated *after* conditioning on finding living individuals of that particular age. Consequently, it automatically assumes that such an individual will indeed be found. In a finite population, sampling issues may make this difficult, since survivorship also strongly declines with age (colour gradient in Fig 3). This means that a large value of *δ*_*µ*_ does not guarantee that deviations from Gompertz are easy to find: the expected survivorship can become very low before *δ*_*µ*_ becomes appreciably large. One way to visualise when deviations become difficult to detect due to low expected survivorship is to calculate *S*(*t*) at the time when the relative deviation *δ*_*µ*_( *f*_0_, *t*)/E[*µ*(f_*t*_)|f_0_ = *f*_0_] ∈ [0, 1) first exceeds a fixed ‘detection threshold’ *θ*. A low *θ* implies that the empirical data admits sufficient statistical power to detect small deviations from Gompertz-Makeham dynamics, whereas setting *θ* higher means that we need a larger difference in the relative mortality rate predictions *δ*_*µ*_/E[*µ*(f_*t*_)|f_0_ = *f*_0_] before we have confidence that the curve being followed is indeed not Gompertz-Makeham and exhibits a detectable plateau (see Camarda (2022) for simulations illustrating how confidence in determining a plateau vs exponential mortality hazard can vary).

In Fig 4, we use the exact expression Eq. 8 to plot the expected survivorship *S* at the age when *δ*_*µ*_/E[*µ*(f_*t*_)|f_0_ = *f*_0_] first exceeds various threshold values *θ*. As the total number of sub-systems *N* increases, the survivorship at the point where a fixed threshold *θ* is exceeded quickly becomes extremely small. Unlike the upper bound Ineq. 10, the expected survivorship (obviously) decreases as extrinsic mortality is increased, causing deviations from Gompertz-Makeham to be difficult to detect (compare Fig 4A vs Fig 4B), despite the deviations being less constrained by Ineq. 10. Loss of expected survivorship is thus an additional way in which observed mortality curves in data can resemble the trajectory of the Tithonus model.

**Figure 4:**
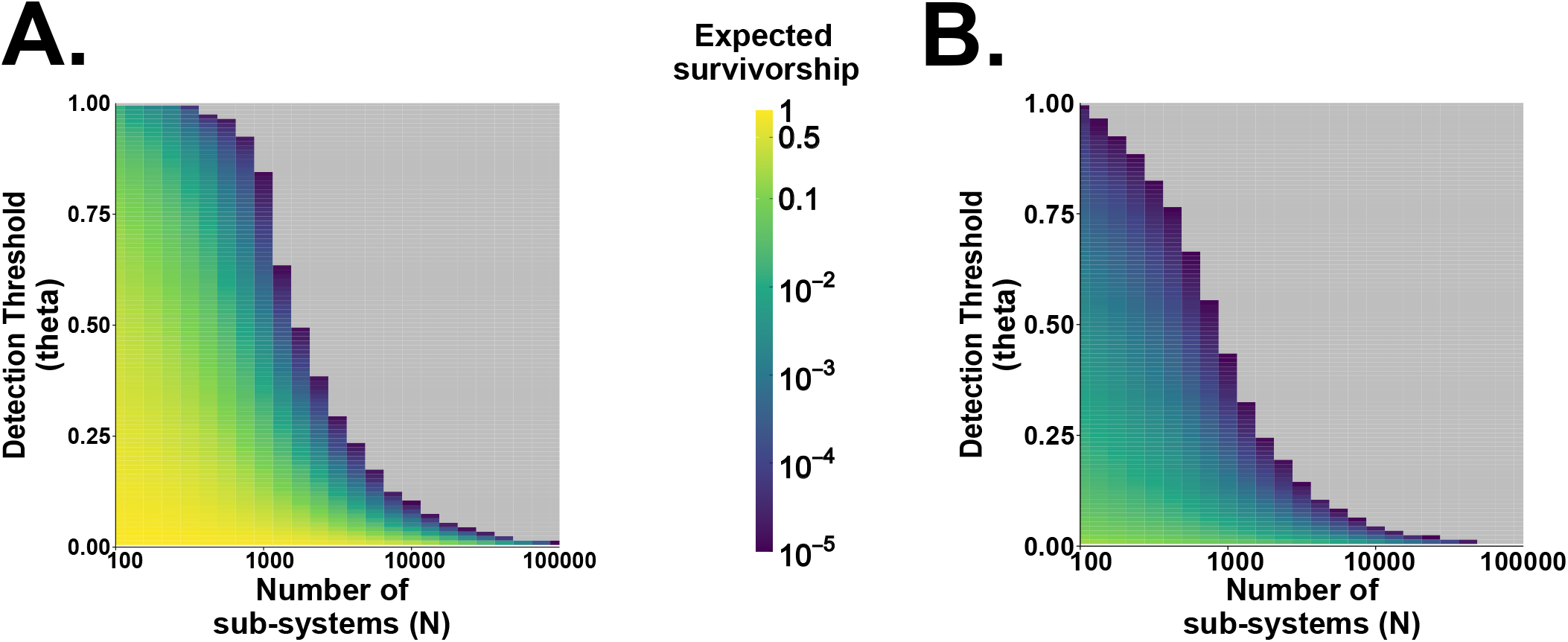
The mean survivorship at the point when the deviation between our complete model and the Tithonus model first exceeds a fixed detectability threshold *θ*. X-axis is the number of sub-systems *N*, and colour is the expected survivorship *S*(*t*) from Eq. 7 at the age *t* when the relative deviation *δ*_*µ*_( *f*_0_, *t*)/E[*µ*(f_*t*_)|f_0_ = *f*_0_] first exceeds the corresponding point on the Y-axis (*θ*). For instance, at *N* = 1000 and *µ*_*e*_ = 10^−5^ (panel A), a 50% deviation of our complete model from the Tithonus model (and in this case Gompertz-Makeham mortality) requires a survivorship of about 10^−3^. Grey colour indicates that either the expected survivorship dropped below 10^−5^ or the detectability threshold *θ* was never reached by *t* = 500. Results are plotted for both **(A) Low extrinsic mortality**, *µ*_*e*_ = 10^−5^, and **(B) High extrinsic mortality**, *µ*_*e*_ = 0.1. Relative deviation *δ*_*µ*_/E[*µ*(f_*t*_)|f_0_ = *f*_0_] and expected survivorship 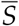 are both calculated using the Feynman-Kac representations by numerically simulating 20,000 realisations of the Tithonus SDEs using the Euler-Maruyama method for each value of *N*. We use *r*_+_( *f* ) = (1 − *f* )(2*ϕ*_*ρ*_ + *k*_*ρ*_ *f* ) and *r*_−_( *f* ) = (1 − *f* )*ϕ*_*ρ*_, so that the failure accumulation rule is *r*( *f* ) = (1 − *f* )(*ϕ*_*ρ*_ + *k*_*ρ*_ *f* ) and the turnover is *τ*( *f* ) = (1 − *f* )(3*ϕ*_*ρ*_ + *k*_*ρ*_ *f* ). We use the mortality rule *µ*( *f* ) = *µ*_*e*_ + *χ f* /(1 − *f* ). Fixed parameter values across both panels: *ϕ*_*ρ*_ = 10^−3^, *k*_*ρ*_ = 0.1, *χ* = 0.2, *f*_0_ = 0.01. We illustrate the effects of varying *k*_*ρ*_ and *χ* in supplementary figure S2.

Undetectability of deviations due to loss of expected survivorship is fundamentally a function of limits on measurement precision and statistical power. Darker hues in 4 indicate that only very few individuals are still alive when deviations of a specific size become visible. Yet, in principle, if one could start with sufficiently many sufficiently (and possibly unrealistically) large cohorts, patterns in data could be detected up to arbitrarily advanced ages (with arbitrarily low survivorship). In contrast, the upper bound Ineq. 9 is a mathematical constraint on the maximum possible value of *δ*_*µ*_; it is hence conservative (*i.e*. the realised deviation could be well below this bound) but unaffected by considerations of sample size or statistical power. Put differently, the expected survivorship 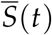 (Eq. 7, colour in Fig 4) tells us the probability of finding a living individual of age *t* in our complete model, while the bound Ineq. 9 tells us how different the typical individual that has reached age *t* (without having died) in our complete model is from the typical Tithonus individual of the same age.

### Biological principles for selecting failure accumulation and mortality rules

In this section, we postulate some desirable properties of biologically reasonable failure accumulation rules (*r*) and mortality rules (*µ*). We then show that the simplest null model fulfilling these properties produces Gompertz-Makeham curves. We also present general scaling laws for any mortality rule which predicts Gompertz-Makeham curves. For simplicity, we focus in this section on the *N* → ∞ limit of our Tithonus model (Eq. 1 and Eq. 2 with *N* → ∞). The modifications to this section induced by selective disappearance and *N* < ∞ can be understood via our complete model using Eq. 6 (figures 2C, 2D, 3, and 4).

We begin with the failure accumulation rule *r*( *f* ). Since the failure rate is assumed to depend on the proportion of failed sub-systems irrespective of their identity, we can define a per-capita failure rate for each functioning sub-system. Since the fraction of functioning sub-systems is 1 − *f*, a sensible general choice for the *r*( *f* ) is thus *r*( *f* ) = (1 − *f* )*ρ*( *f* ), where the function *ρ*( *f* ) > 0, returns the failure rate of each functioning sub-system when a proportion *f* have failed. We call its partial derivative *ρ*^′^( *f* ) ‘interdependence’: it quantifies how the per-capita failure rate changes as the proportion of failed sub-systems changes. We capture ‘Failure begets failure’ dynamics ( also called ‘vicious cycles’ of dysfunction (Belikov, 2019), see also (Tian et al., 2023)) by demanding that *ρ*^′^( *f* ) > 0. Large *ρ*^′^( *f* ) applies to extremely integrated/unitary organisms with many complex interdependencies, while low values indicate modularity in which subsystems continue to be functional even if their neighbours fail or die. We have

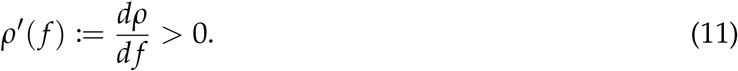

Reasonable choices for the failure rule *µ*( *f* ) must have *µ*( *f* ) ≥ 0 (optionally, we may also set *µ*(0) > 0 when including extrinsic causes of mortality that arise even in the absence of failures), and all else being equal, individuals with more failures should have a higher mortality rate: *µ*^′^( *f* ) > 0. An organism should also not remain alive when all of its sub-systems have failed, therefore 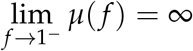 which guarantees instantaneous death (infinite mortality hazard) at the point *f* = 1. As we will see in the following sub-sections, Gompertz-Makeham mortality is tied to the rate at which *µ* approaches infinity as *f* → 1.

#### A simple null: logistic failure accumulation and geometric mortality produces Gompertz-Makeham mortality curves

During the initial part of the failure accumulation process (small *f* − *f*_0_), a Taylor expansion for the per-capita failure rate yields *ρ*(*f*)≈*ϕ*_*ρ*_ + *k*_*ρ*_*f*, defining 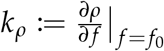 and *ϕ*_*ρ*_ := *ρ*_0_ − *k*_*ρ*_ *f*_0_. This implies

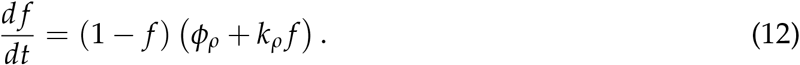

for the failure dynamics. We retain the *ρ* in the subscripts as a reminder that *ϕ*_*ρ*_ and *k*_*ρ*_ are defined *for a given per-capita failure rate function ρ*( *f* ), but henceforth assume the linearisation holds exactly henceforth (we discuss relaxing this assumption in the next subsection). Eq. 12 is logistic-like (it is the logistic equation when *ϕ*_*ρ*_ = 0) and can be solved exactly using partial fractions.

Since we required *r*(0) > 0, Eq. 12 implies *ϕ*_*ρ*_ > 0. The interdependence *ρ*^′^( *f* ) = *k*_*ρ*_ is a constant. From Eq. 11 it follows that *k*_*ρ*_ is non-negative. With *ϕ*_*ρ*_ and *k*_*ρ*_ both non-negative, the functional form Eq. 12 admits a biologically appealing interpretation: The parameter *ϕ*_*ρ*_ acts as a constant baseline failure rate of each sub-system (a measure of the resilience of each individual sub-system; it could be affected by external parameters such as hazardous environments), and *k*_*ρ*_ represents the ‘failure begets failure’ dynamics: it is a measure of the integratedness of the organism as a whole. See the supplementary section S6 for an alternative derivation of Eq. 12 from a mechanistic network-structured model.

What is a simple null mortality rule? A linear function such as *µ*( *f* ) = *µ*_*e*_ + *χ f* (as suggested e.g. by Gavrilov and Gavrilova (1991)) does not satisfy our requirement that complete failure leads to instantaneous death. A simple mortality rule that satisfies all the desirable properties we mentioned is what we call a ‘geometric mortality rule’

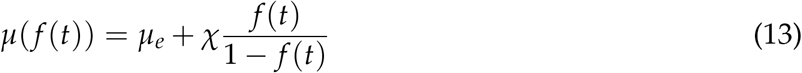

Once again, the first and second term of the RHS of Eq. 13 capture extrinsic mortality and the effect of failed subsystems, respectively. Eq. 13 can be written as an infinite geometric series *µ*(( *f* (*t*)) = *µ*_*e*_ + *χ f* (*t*) + *f* ^2^(*t*) + *f* ^3^(*t*) + *f* ^4^(*t*) + · · ·, providing intuition for the failure-dependent part Eq. 13 as the simplest choice that reflects the need for a linear effect of failures on mortality, a quadratic effect due to synergies between pairs of sub-systems, a cubic effect due to synergies between triads of sub-systems, etc. Note that we cannot stop this logic at any finite *f*^*k*^, because doing so would make *µ*(1) = *µ*_*e*_ + *χ*^*k*^ finite (whereas we required 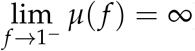) Substituting the solution of Eq. 12 into Eq. 13 yields, after some algebra,

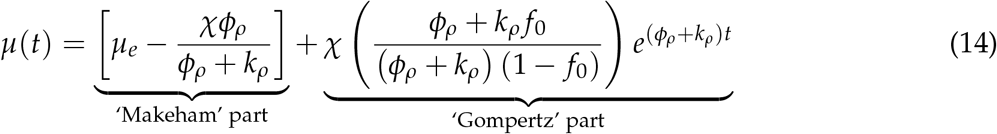

Here, logistic failure accumulation (Eq. 12) together with the geometric mortality rule (Eq. 13) leads to (exact) Gompertz-Makeham mortality curves in the *N* → ∞ Tithonus model (see Fig 2 for the case when *N* < ∞). In our complete model, there is a late-life mortality plateau due to selective disappearance whenever *N* < ∞ (Fig 3), but this may be difficult to detect in data (Fig 4).

#### Beyond the simple null: Gompertz-Makeham is tied to inverse power-law singularities of the mortality rule

Ideally, the failure accumulation process within an organism should be derived from a concrete, underlying mechanistic process, using data-driven models (Farrell et al., 2021) or reliability theory (Gavrilov and Gavrilova, 2001; Boonekamp et al., 2015). As a result, however, the failure accumulation may become very complex, and the linearisation we applied above may be a poor approximation (especially as *f* − *f*_0_ grows). For failure rules that are higher order than Eq. 12, the geometric rule Eq. 13 will not generally lead to exact Gompertz-Makeham dynamics. Equally, even if we assume the logistic failure rule Eq. 12 holds exactly, it is possible to construct mortality rules do not lead to Gompertz-Makeham mortality despite satisfying our biological criteria (e.g. consider *µ*( *f* ) = *µ*_*e*_ − *χ* log(1 − *f* )). Thus, we ask an inverse question: Given a (possibly very complicated) per-capita failure accumulation rate *ρ*( *f* ), which mortality rules generate exponential mortality curves?

In supplementary section S7, we derive a precise mathematical relation (Eq. S75) between the failure accumulation rule, the given Gompertz-Makeham parameters, and the mortality rule that is both necessary and sufficient to generate Gompertz-Makeham curves of known parameter values in the *N* → ∞ Tithonus model. If the Gompertz-Makeham parameters are not known, Eq. S75 provides a two-parameter family of mortality rules that yield exact Gompertz-Makeham curves in this model. When *ρ*(1) < ∞, Eq. S75 implies that a mortality rule must necessarily scale as *µ*( *f* ) ∼ (1 − *f* )^−*β*/*ρ*(1)^ near complete failure ( *f* = 1) to yield Gompertz-Makeham curves. Here, *β* is the rate parameter of the emergent Gompertz-Makeham curve. While this is a concrete, testable prediction, estimating the mortality rates of organisms near complete system failure ( *f* = 1) will require exceptional experimental ingenuity.

In supplementary section S7, we also demonstrate that if the mortality rule exhibits an inverse power law singularity of the form (1 − *f* )^−*γ*^ near complete failure ( *f* = 1) for *some* scaling exponent *γ* > 0 (Eq. S84), then the overall mortality curve follows a Gompertz curve asymptotically (i.e. after a potential early life transient deviation) with rate parameter *γρ*(1) for any increasing (*ρ*^′^( *f* ) > 0) and bounded (*ρ*(1) < ∞) failure accumulation rule.

The asymptotic sufficiency result provides an additional rationale for viewing the geometric mortality rule Eq. 13 as a simple null. It has the scaling exponent *γ* = 1 and hence provides the simplest example that produces Gompertz behaviour asymptotically (i.e. at late ages). In particular, any increasing (*ρ*^′^ > 0) and bounded (*ρ*(1) < ∞) failure accumulation rule can be coupled with the geometric mortality rule, and the result will asymptotically become a Gompertz curve with rate parameter *ρ*(1). Note, however, that the non-Gompertzian transient may be long, and our complete model shows that selective disappearance may cause deviations at the population level later in life. Given an arbitrary *ρ* and *µ* with *µ*( *f* ) ∼ (1 − *f* )^−*γ*^ as *f* → 1, it is thus possible that early-life deviations (transient non-Gompertzian dynamics) and late-life deviations (deviations due to selective disappearance) overlap, causing the entire population-level mortality curve to be non-Gompertzian.

Lastly, supplementary section S7.3 demonstrates that greater ‘interdependence’ *ρ*^′^( *f* ) will lead to a faster rate of ageing *ceteris paribus* whenever the overall mortality curve is either exactly or asymptotically Gompertz-Makeham. The logistic failure accumulation rule Eq. 12 has *ρ*^′^( *f* ) = *k*_*ρ*_; accordingly, larger *k*_*ρ*_ leads to a faster rate of ageing.

## Discussion

George Williams, one of the pioneers of senescence theory, was quite pessimistic about both reliability theoretic thinking and Gompertz-Makeham curves towards the end of his life (Williams, 1999), even referring to the same Tithonus myth we use here (though he used the metaphor to criticise a focus on lifespan shortening at the expense of senescent processes of individuals that are still alive). Williams’ worries included that the ‘organisms as failing machines’ analogy could not handle repair or replacement of failed systems, that Gompertz-Makeham mortality was studied phenomenologically via statistical fitting as opposed to mechanistic understanding, and the pattern was not robust against heterogeneity. In this manuscript, we have provided a general theoretical framework of senescence arising from stochastic failures of (or damage to) intra-organismal sub-systems that counters all these criticisms. Firstly, our framework incorporates repair, only assuming that repair cannot perfectly balance failures forever, not least because repair mechanisms themselves can fail (Gorbunova et al., 2007). Secondly, we derive exponential mortality curves as the null result of a model arising from two mechanistically grounded observations: (i) organisms are integrated systems with many interdependencies and thus failure begets failure, and (ii) death must be immediate if all of an organism’s sub-systems have failed. Thirdly, we demonstrate that while (stochastic) heterogeneity does cause deviations from Gompertz-Makeham curves (also see Weitz and Fraser, 2001), these deviations may often be so small as to be undetectable in data.

Importantly, our models assume every individual is born with identical parameters and the same initial condition *f*_0_, and inter-individual differences emerge only thereafter, in the absence of any intrinsic differences in ‘quality’. These differences in biological age between individuals of the same chronological age fundamentally emerges due to stochasticity in the failure accumulation process, and individuals who (stochastically) accumulate less damage become overrepresented at later ages. Our work hence adds to growing literature on the importance of the observation that stochastic variation accumulates with age (López-Otín et al., 2023; Meyer and Schumacher, 2024; the 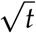 in Eq. 10). Empirical evidence likewise points to age-related dysfunctions having a strong stochastic component (López-Otín et al., 2023; Meyer and Schumacher, 2024; Meyer et al., 2025). Our work demonstrates how such stochasticity can be incorporated into mechanistic models of senescence from the ground up using birth-death processes and diffusion approximations. Our framework hence bridges the gap between mechanistic reliability theoretic style models of damage accumulation and random-walk models of senescence such as Woodbury and Manton (1983) and Yashin et al. (1985) (see Supplementary section S8). The observation that stochasticity generates dynamic variation between initially identical individuals has also been made in contexts unrelated to senescence, see Tuljapurkar et al.’s (2009) ‘dynamic heterogeneity’ and Snyder et al.’s (2021) ‘state trajectory luck’.

Our results are aligned with Weitz and Fraser (2001) and Ledberg (2020). The former work explores the idea that stochasticity yields mortality plateaus, with a model that is similar to ours but less realistic in some key assumptions (see supplementary section S8). The latter work showed that a broad class of models of stochastic damage accumulation predict (approximately) exponential mortality curves, albeit with a model structure that is quite different to ours. Ledberg (2020) assumes mortality is proportional to the probability that the number of failed sub-systems ‘queuing to be repaired’ exceeds a fixed threshold, while we assume mortality is a continuous function of the proportion of failed sub-systems. The fact that these two very different approaches predict similar outcomes further strengthens the view that the prediction of exponential mortality curves is often robust to the gory mechanistic details of the wear and tear that accumulates in life.

Some of our work’s novelty is based on avoiding particular sub-system interdependency structures (‘series’, ‘parallel’, ‘scale-free’, etc.) in order to understand the qualitative behaviour of mortality curves. Of course, assessing specific quantitative aspects may require deeper structure (for examples see Vural et al., 2014; Sun et al., 2020). Our main approach instead outlines general principles that biologically reasonable failure accumulation and mortality rules should obey, and explores the consequences. This provides biological justification for previous abstract state space models (Woodbury and Manton, 1977; Yashin et al., 1985) (for details see supplementary section S8). The simplest failure accumulation rule (‘logistic’, Eq. 12) and mortality rule (‘geometric’, Eq. 13) that satisfy our proposed principles predict exact Gompertz-Makeham mortality curves in our Tithonus model, providing a null model for exponential mortality and late-life plateaus when populations consists of individuals with stochastic failure accumulation with dynamic heterogeneity. Going beyond the null model, many failure accumulation rules are approximated by the logistic failure rule in the initial part of the failure accumulation process (*i.e*. early in life, when the Taylor expansion in Eq. 12 remains accurate); in late life, in turn, many mortality rules become asymptotically Gompertz. Furthermore, for each failure accumulation rule, there is a two-parameter family of mortality rules that lead to exact Gompertz-Makeham mortality (Eq. S75).

Though our general approach is agnostic to the precise details of the interdependencies between sub-systems, we also sketch a brief outline of how to incorporate more precise structure in supplementary section S6. There, we use a model inspired by Nielsen et al. (2024) to derive an approximate failure rule for organisms whose sub-systems are organised as vertex-transitive graphs (networks) and the failure rate of a sub-system depends linearly on its number of failed neighbours. We show that the rate of the emergent Gompertz-Makeham curve is controlled by the degree of the graph, and, interestingly, find that this dependence disappears if the failure rate of a sub-system depends on the *proportion* of failed neighbours rather than the *number* of failed neighbours. However, our toy model in the supplementary is only valid for a class of graphs in which every vertex has the same degree. Since we biologically expect some sub-systems to be more central than others, extensions to study the effect of heterogeneity in the degree distribution are valuable, especially given recent work in network science highlighting the stabilising nature of such heterogeneity (Montanari et al., 2026). Heterogeneity between sub-systems could also manifest in terms of directional dependencies in function, formally modelled via digraphs. Directionality becomes important in ‘multi-stage’ failure cascades (Armitage and Doll, 1954; Webster, 2019), and extending our framework to account for this possibility may prove fruitful, with Webster (2019) offering a promising starting point.

### Mortality plateaus and statistical predictions

Our work touches on an important, if somewhat overlooked, theoretical conundrum relevant to humans and related unitary organisms: if selective disappearance is inevitable and always lowers late life mortality (thus causing a plateau), why do so many organisms exhibit an excellent fit of the Gompertzian model to data? In our model, stochasticity creates heterogeneity between individuals of the same chronological age, and this heterogeneity can cause late-life plateaus due to selective disappearance. However, predicted deviations may often be too small to detect in data, and we shoaw that deviations are smaller when when the causal route to each intrinsic death is distinct, *i.e*. different organisms are expected to have had a different sequence of sub-system failures before mortality finally strikes. This is relatable for anyone studying human causes of death, which — despite attempts to bin them into broad-brush categories such as ‘cardiovascular issues’ and ‘cancer’ — are extremely diverse. For humans in particular, therefore, our modelling framework suggests that deviations from Gompertz-Makeham curves should generally be negligible.

Deviations from the Tithonus model (and thus Gompertz-Makeham curves) may also be undetectable in many biologically relevant cases if expected survivorship becomes small before the deviation can become large (background gradient in Fig 4). In other words, if most lives end while Tithonus still holds well, it is statistically difficult to prove that the deviation exists unless one has very large cohorts. This appears to be the case for humans, where the very existence of late-life plateaus is contentious. The debate focuses on centenarians and beyond, a cohort where most deaths have already occurred (Barbi et al., 2018; Feehan, 2018; Newman, 2018). Camarda (2022) has used simulations to show that statistical tests which assume a mortality plateau as the null typically lack the statistical power to detect significant deviations from the null in the oldest old.

Mortality plateaus exist in a documentable manner in a few well-studied invertebrates (table 1). This is in line with our complete model that shows (a) plateaus should generally exist, in the sense that the deviation Eq. 8 is exactly zero only under infinite *N* or if the measurement does not covary with survivorship, and (b) plateaus should not always remain undetectable and should be more readily detectable with higher sample sizes and fewer distinct pathways to death. Indeed, studies that have found plateaus in invertebrates typically use very well-controlled experimental conditions that minimise the number of distinct ways to die and further involve large sample sizes, typically between 10^3^ − 10^6^ (see table 1).

**Table 1.** Empirical studies finding late-life mortality plateaus in non-human animals. ‘Cohort size’ is the initial cohort size at the beginning of the experiment/observation. ‘gen. hom. cohort’ means that the initial cohort was generated from a highly inbred or clonal line to minimise the possibilities of unobserved heterogeneity due to genetic differences (fixed frailty classes). When sex-specific curves are reported, we present both sample sizes. We exclude human studies because a table does not give justice to the complexities of the statistical issues involved (see main text and also Barbi et al. (2018), Feehan (2018), Newman (2018), Gavrilov and Gavrilova (2019), and Camarda (2022)). Table 1 in Pletcher and Curtsinger (1998) shows a similar table reviewing empirical demonstration of plateaus from 13 different stocks of *Drosophila melanogaster* (sample sizes of the studies collated in Pletcher and Curtsinger (1998) range from 2328 to 109,860).

| **Species** | **Taxon** | **Cohort size** | **Reference** |
|---|---|---|---|
| *Musca domestica* | Diptera (house fly) | 4650 | Gavrilov and Gavrilova, 2006 (Fig 5.5) |
| *Drosophila melanogaster* | Diptera (fruit fly) | 5751 | Curtsinger et al., 1992 |
| *D. melanogaster* (gen. hom. cohort) | Diptera (fruit fly) | 6338 | Vaupel et al., 1998 |
| *Ceratitis capitata* | Diptera (med fly) | 1,203,646 | Carey et al., 1992 |
| *Anastrepha serpentina* | Diptera (fruit fly) | 341,134 | Vaupel et al., 1998 |
| *Anastrepha obliqua* | Diptera (fruit fly) | 297,087 | Vaupel et al., 1998 |
| *Anastrepha ludens* | Diptera (fruit fly) | 851,100 | Vaupel et al., 1998 |
| *Diachasmimorpha longiacaudata* | Hymenoptera (parasitoid wasp) | 27,542 | Vaupel et al., 1998 |
| *Callosobruchus maculatus* | Coleoptera (seed beetle) | 4299 (F), 4395 (M) | Fox and Moya-Laraño, 2003 (Fig 2A) |
| *Stator limbatus* | Coleoptera (seed beetle) | 1411 (F), 1401 (M) | Fox and Moya-Laraño, 2003 (Fig 3A) |
| *C. elegans* (gen. hom. cohort) | Nematode | ≈ 550,000 | Vaupel et al., 1998 |

Previous work has shown that mortality plateaus may arise as a logical inevitability in broad classes of models (Fox and Moya-Laraño, 2003; Steinsaltz and Evans, 2004) and that radically different biological hypotheses for ageing can be statistically indistinguishable given mortality curve data alone (Yashin et al., 1994; Camarda, 2022). Consequently, determining the causalities that underlie an observed mortality curve is empirically challenging. Work such as Yashin et al. (1994) suggests that it can be very difficult to make informative statements about biological processes at the individual level (e.g. distinguishing between programmed ageing theories vs damage accumulation theories) by testing the goodness-of-fit of census data to the predicted population-level mortality curve in the absence of a more mechanistic understanding of the individual-level (failure accumulation) process. However, our results provide an alternative route to falsifying our framework if a candidate failure accumulation rule is identified and Gompertz-Makeham is assumed to be a good description of population level mortality.

In particular, if we know (or guess) the functional form of the per-capita failure rate *ρ*( *f* ) of a sub-system on mechanistic grounds such as through reliability theory (Gavrilov and Gavrilova, 2001) or data-driven model specification (Farrell et al., 2021), an inverse power-law scaling *µ*( *f* ) ∼ (1 − *f* )^−*β*/*ρ*(1)^ near *f* = 1 for a constant *β* > 0 is a *necessary* requirement for Gompertz-Makeham curves to emerge in our framework (supplementary section S7). This means that when the proportion of already failed sub-systems is very high, we predict that the regression of log(*µ*) against log(1 − *f* ) is a straight line with slope proportional to −1/*ρ*(1). We also show (supplementary section S7) that a scaling of the form *µ*( *f* ) ∼ (1 − *f* )^−*γ*^ near *f* = 1 for any *γ* > 0 is *sufficient* to generate mortality curves that asymptotically converge to Gompertz curves later in life in our framework. However, note that late-life mortality patterns are often affected by selective disappearance, as illustrated by our complete model (Fig 3).

### Not all organisms are unitary

Our results are fundamentally impacted by the ‘failure begets failure’ idea, but the ‘begetting’ does not necessarily operate equally strongly in all taxa. Finch (1994) has hypothesised that the potential for continual replacement of damaged modules in modular and non-unitary organisms (e.g. colonial ascidians, corals, many vascular plants) should yield a lower rate of senescence, an idea that has received some tentative empirical support (Finch, 2009; Bernard et al., 2020). In our model, the interdependence *ρ*^′^( *f* ) is a measure of organismal integratedness, and supplementary section S7.3 shows that individuals with greater interdependence will exhibit a greater rate of senescence *ceteris paribus*. Our work thus lends theoretical support to Finch’s hypothesis. In supplementary section S6, we also formulate an explicit graph (network) structured failure accumulation rule (see also Nielsen et al. (2024)) and demonstrate that the rate parameter of the resultant Gompertz-Makeham curve is controlled by the degree of the graph (the number of sub-systems that each sub-system depends on).

Modular organisms are often also mentioned as excellent candidates for showing not just slowing down of senescence, but making it negligible or even negative (Bernard et al., 2020). Our present models preclude this possibility (Baudisch, 2012) by assuming the expected failure accumulation rate *r*( *f* ) is always positive and increasing in *f* . Though our approach of modelling failure accumulation as (directed) diffusion with killing does not require this assumption, our results on Gompertz curves do. Future work could consider failure accumulation rules that permit an intermediate value *f* ^∗^ ∈ (0, 1) at which *r*_+_( *f* ^∗^) = *r*_−_( *f* ^∗^). This creates the possibility of an internal equilibrium where failure is exactly balanced by repair, leading to more complex mortality patterns than considered here, potentially including negative or negligible senescence. Biologically, however, it may be unrealistic to expect failures to be balanced by repair forever (repair mechanisms may themselves fail (Gorbunova et al., 2007)).

### Extensions and Outlook

Our manuscript has focused on actuarial rather than reproductive senescence for the sake of concision, despite many empirical studies on selective disappearance focusing on the latter (Bouwhuis et al., 2009; Hayward et al., 2013; Hämäläinen et al., 2014). Since the ‘measurement’ in equations such as Eq. 8 can be any nice function, it could easily be used to model reproductive output and related notions of fecundity as long as the quantity of interest only depends on chronological age through the failures acquired by that age. For instance, if failures cause a linear decline in reproductive output, we can model reproductive senescence via the measurement function *M*( *f* ) = *m*_max_ − *λf* for constants *m*_max_, *λ*, and it is easily verified that though *M* decreases with age, selective disappearance causes an apparent increase very late in life, generating apparent terminal effort (this is of course the reproductive analog of late-life mortality plateaus). Deriving general results about reproductive senescence systematically is beyond the current scope of our paper, but provides a potentially fruitful research avenue.

We have also assumed that there is no variation in the state of newborns. We chose this to underscore that selective disappearance can operate without fixed frailty classes, but mixture distribution theory (Finkelstein and Esaulova, 2006) can be used to extend our framework to populations where such classes exist. Since fixed frailty classes cause average mortality to be ‘pulled down’ as less frail individuals are overrepresented at advanced ages (Finkelstein and Esaulova, 2006; Vaupel and Missov, 2014), we conjecture that adding fixed frailty classes to our stochastic model produce Siler-like curves (Siler, 1979).

Reproduction and survival, of course, may be linked via trade-offs, and both may further trade off with growth (Vaupel et al., 2004; Baudisch, 2012; Maklakov and Chapman, 2019; Cohen et al., 2020; Avila and Lehmann, 2023; Chmilar et al., 2024). A multi-dimensional (vector-valued) *f* could conceptually represent sub-systems that contribute to distinct organismal functions such as fecundity and survival, with trade-offs modelled via stochastic optimal control theory (Avila et al., 2021; Avila and Lehmann, 2023). The straightforward multi-dimensional extension of our logistic failure accumulation rule resembles a generalised Lotka-Volterra equation, and thus tools such as random matrix theory that have found widespread application in mathematical community ecology (Allesina, 2026) could be fruitfully imported our framework.

We have not modelled evolutionary dynamics, but the realised expected path through failure space is of course subject to evolutionary forces and thus our model could also be extended to the evolutionary setting. We thus finish by commenting on key papers that have put damage or failure accumulation into an evolutionary framework. In Laird and Sherratt (2009), individuals consist of multiple fully redundant (‘in parallel’) sub-systems, and the number of such sub-systems evolves; and Laird and Sherratt, 2010 considered the same when sub-systems are arranged in a ‘cascade’. Cast in our language, these models hold the failure rule, mortality rule, and interdependence structure constant while evolution impacts the total number of sub-systems (which we kept constant within each scenario). Bega and Hadany (2026), in turn, considered how failure accumulation models interact with the selection shadow Hamilton’s (1966) and its evolutionary consequences. Though Bega and Hadany (2026) present a conceptual starting point to studying how parameters like our *k*_*ρ*_ and *ϕ*_*ρ*_ evolve and affect the evolution of lifespan, they make an unfortunate assumption that Gompertz parameters can vary independently of each other (for why this is problematic, see (Strehler and Mildvan, 1960) and our supplementary section S8). Merging our framework with Laird and Sherratt (2009, 2010) and Bega and Hadany (2026), with some of the issues fixed, could allow progress towards unified understanding of how selective disappearance, interdependencies between sub-systems, and mortality due to failure accumulation together determine the ultimate evolutionary patterns of senescence. Ideally, such a model will also take eco-evolutionary aspects seriously by incorporating density dependence, as it is known to impact the evolution of senescence when extrinsic mortality can kill individuals that are not yet senescent (de Vries et al., 2023).

Lastly, it has not escaped our notice that Eq. 4 bears a striking resemblance to equations arising in the multi-level selection literature (Kimura, 1984, Eq. 2.5; Fontanari and Serva, 2014, Eq. 11; Luo, 2014, Eq. 1; Cooney et al., 2023, Eq. 5). The conceptual connections between senescence and multi-level selection suggested by this similarity are made precise in Bhat and Kokko (submitted).

## Data and Code Availability

All code and generated data used to generate the figures within this manuscript are available at https://anonymous.4open.science/r/ageing_failures_scripts/.

## Appendix A: Summary of recurring notation

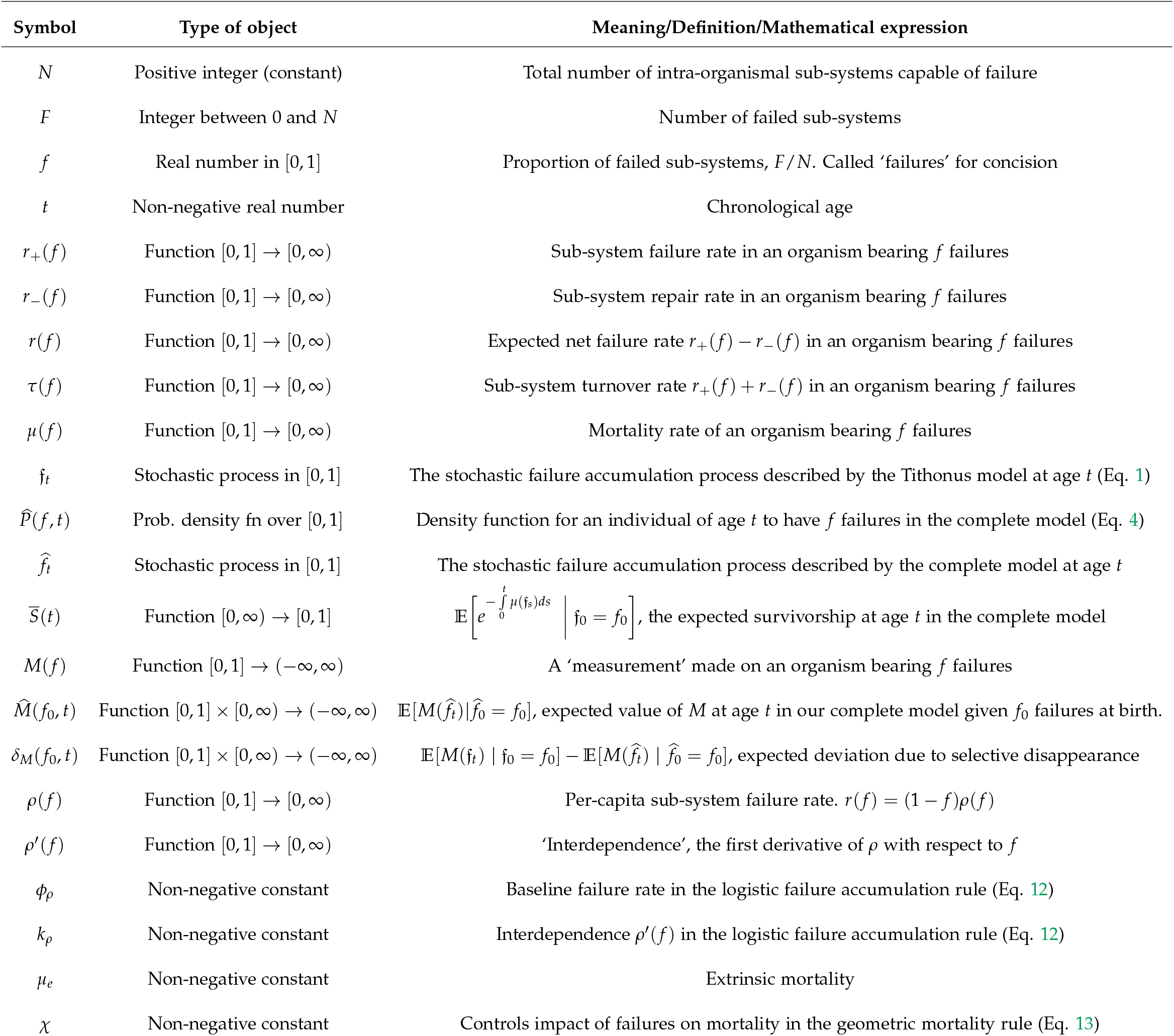

| Symbol | Type of object | Meaning/Definition/Mathematical expression |
|---|---|---|
| *N* | Positive integer (constant) | Total number of intra-organismal sub-systems capable of failure |
| *F* | Integer between 0 and *N* | Number of failed sub-systems |
| *f* | Real number in [0, 1] | Proportion of failed sub-systems, *F*/*N*. Called ‘failures’ for concision |
| *t* | Non-negative real number | Chronological age |
| *r*<sub>+</sub>(*f*) | Function [0, 1] → [0, ∞) | Sub-system failure rate in an organism bearing *f* failures |
| *r*<sub>−</sub>(*f*) | Function [0, 1] → [0, ∞) | Sub-system repair rate in an organism bearing *f* failures |
| *r*(*f*) | Function [0, 1] → [0, ∞) | Expected net failure rate *r*<sub>+</sub>(*f*) − *r*<sub>−</sub>(*f*) in an organism bearing *f* failures |
| *τ*(*f*) | Function [0, 1] → [0, ∞) | Sub-system turnover rate *r*<sub>+</sub>(*f*) + *r*<sub>−</sub>(*f*) in an organism bearing *f* failures |
| *μ*(*f*) | Function [0, 1] → [0, ∞) | Mortality rate of an organism bearing *f* failures |
| 𝔣<sub>*t*</sub> | Stochastic process in [0, 1] | The stochastic failure accumulation process described by the Tithonus model at age *t* (Eq. 1) |
| $\widehat{P}(f,t)$ | Prob. density fn over [0, 1] | Density function for an individual of age *t* to have *f* failures in the complete model (Eq. 4) |
| $\widehat{f}_t$ | Stochastic process in [0, 1] | The stochastic failure accumulation process described by the complete model at age *t* |
| $\overline{S}(t)$ | Function [0, ∞) → [0, 1] | \$\mathbb{E}\left[e^{-\int_0^t \mu(\mathfrak{f}_s)ds} \,\middle | \, \mathfrak{f}_0 = f_0\right]\$, the expected survivorship at age *t* in the complete model |
| *M*(*f*) | Function [0, 1] → (−∞, ∞) | A ‘measurement’ made on an organism bearing *f* failures |
| $\widehat{M}(f_0,t)$ | Function [0, 1] × [0, ∞) → (−∞, ∞) | $\mathbb{E}[M(\widehat{f}_t)\|\widehat{f}_0 = f_0]$, expected value of *M* at age *t* in our complete model given *f*<sub>0</sub> failures at birth. |
| *δ*<sub>*M*</sub>(*f*<sub>0</sub>, *t*) | Function [0, 1] × [0, ∞) → (−∞, ∞) | $\mathbb{E}[M(\mathfrak{f}_t) \mid \mathfrak{f}_0 = f_0] - \mathbb{E}[M(\widehat{f}_t) \mid \widehat{f}_0 = f_0]$, expected deviation due to selective disappearance |
| *ρ*(*f*) | Function [0, 1] → [0, ∞) | Per-capita sub-system failure rate. *r*(*f*) = (1 − *f*)*ρ*(*f*) |
| *ρ*′(*f*) | Function [0, 1] → [0, ∞) | ‘Interdependence’, the first derivative of *ρ* with respect to *f* |
| *ϕ*<sub>*ρ*</sub> | Non-negative constant | Baseline failure rate in the logistic failure accumulation rule (Eq. 12) |
| *k*<sub>*ρ*</sub> | Non-negative constant | Interdependence *ρ*′(*f*) in the logistic failure accumulation rule (Eq. 12) |
| *μ*<sub>*e*</sub> | Non-negative constant | Extrinsic mortality |
| *χ* | Non-negative constant | Controls impact of failures on mortality in the geometric mortality rule (Eq. 13) |

## Supplementary Information

### S1 Deriving the Fokker-Planck-Kolmogorov equations

We present the derivation here for the main model with disappearance. The derivation for the Tithonus model is obtained by setting *µ* ≡ 0 in all calculations in this section.

#### S1.1 Definition and master equation

In the main text, we formulated a continuous time stochastic process for failure accumulation with stochastic mortality. This process took values in 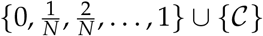, where 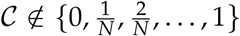 is a ‘cemetery state’. The process is defined by the transitions

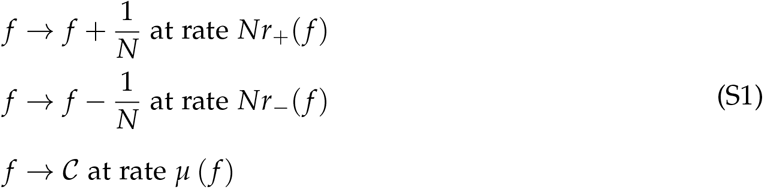

where *r*_±_ are *O*(1) functions. We are interested in computing *P*( *f, t*), the probability that an individual of age *t* has *f* failures, given that individuals are born with *f*_0_ failures at age 0.

To do this, we imagine tracking a large cohort of independent individuals from birth. In a large ensemble of individuals of age *t*, a proportion *P*( *f, t*) of them will have a fraction *f* failures (by definition of probability). We can now simply measure the ‘inflow’ and ‘outflow’ of individuals from each state (Figure S1). Individuals with 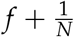 failures flow into the state *f* by repairing a failed sub-system. Repair occurs at rate 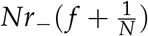 and a fraction 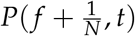 of the cohort consists of such individuals. Similarly, individuals with 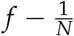 failures flow into the state *f* by accumulating another failure. Failure is at rate 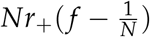 and a fraction 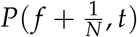 of the cohort consists of such individuals. Thus, the rate of ‘inflow’ to the state *f* is given by

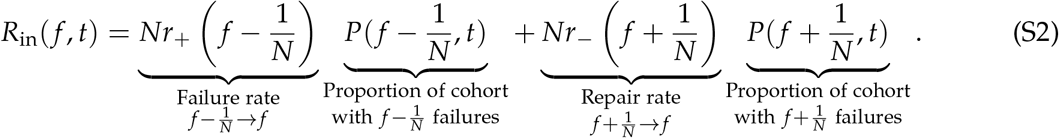

Similarly, an individual with *f* failures may move into a different state by either accumulating another failure, repairing an existing failure, or dying. Thus, the rate of ‘outflow’ from the state *f* is

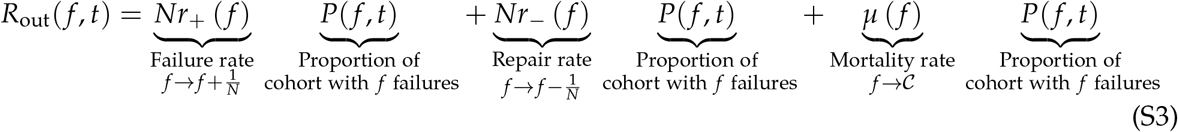

**Figure S1:**
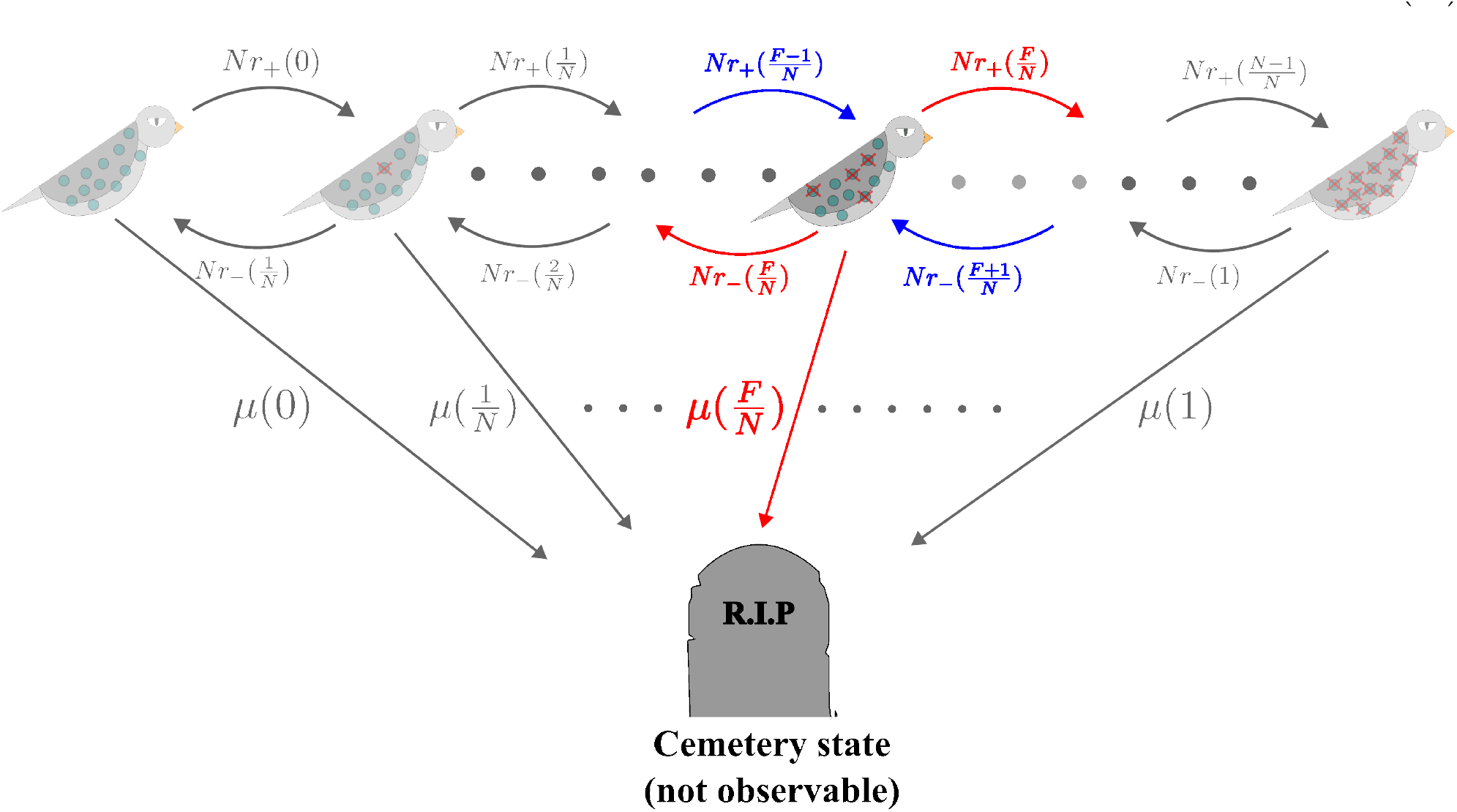
Schematic description of the possible transitions of our process. For a given state *f*, the blue arrows depict the rate of ‘inflow’ to the state, whereas the red arrows depict the rate of ‘outflow’.

The rate of change of *P*( *f, t*) is given by the rate of inflow minus the rate of outflow. Thus, we have

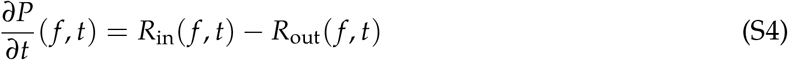

For convenience, let us define two ‘step operators’ 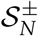, which act on any functions of failures by either adding or removing a failure, *i.e*

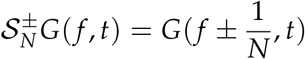

Substituting Eq. S2 and Eq. S3 into the RHS of (S4) and using the step operators, we obtain the compact expression

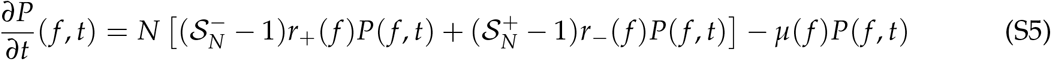

Equation S5 is called a ‘master equation’ or ‘Chapman-Kolmogorov equation’ and completely describes our system exactly. To obtain the expressions presented in the main text, we now carry out a diffusion approximation.

#### S1.2 The diffusion approximation

Since the stochastic process defined by Eq. S1 takes values in 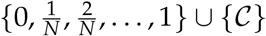 and jumps in units of 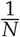, the transitions of *f* within the space of failures 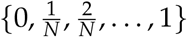 begins to ‘look continuous’ as *N* grows very large. We thus Taylor expand^1^ the action of the step operators 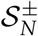 to find

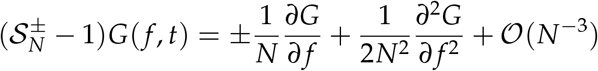

and thus

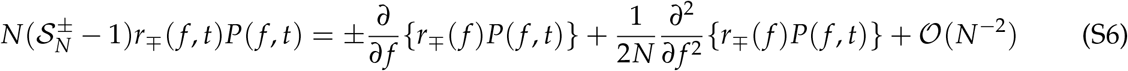

We now perform the diffusion approximation, which consists of neglecting terms of *O*(*N*^−2^). Biologically, neglecting the higher order terms amounts to saying that *f*_*t*_, viewed as a random variable, is entirely characterised by its first two moments, and thus can be thought of as a Gaussian approximation (Black and McKane, 2012). Substituting Eq. S6 into Eq. S5 now yields, after some lines of algebra,

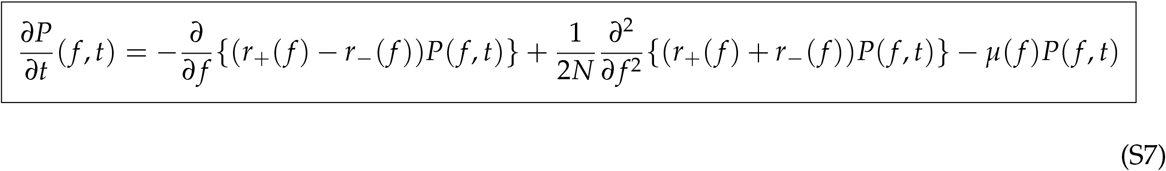

The derivation of the equation for the conditional probability density Eq. 4 starting from Eq. is more involved and appears in Yashin et al., 1985 (their Appendix A), so we do not repeat it here.

### S2 The Feynman-Kac representation of the killed diffusion

In this section, we elaborate on how one arrives at the Feynman-Kac representations presented in the main text. To do this, we first need an alternate characterisation of the stochastic processes we study in terms of their *infinitesimal generators* (Øksendal, 1998, definition 7.3.1; Karatzas and Shreve, 1998, section 5.1). Given a time homogeneous Markov process *X*_*t*_ that lives in a domain *D* ⊆ ℝ, the *infinitesimal generator* of *X*_*t*_ is defined as the (unique) operator *G* satisfying

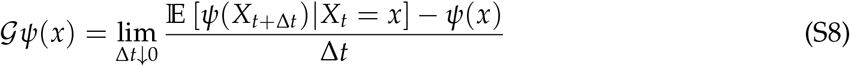

for every ‘nice’^2^ function *ψ* : *D* → ℝ. We require these generators because the Feynman-Kac formula is stated in terms of generators. Rather than asserting the form of the generator of a killed diffusion, we will derive the infinitesimal generator of our stochastic process starting from Eq. S7, thus demonstrating that the process we obtain after our diffusion approximation (system size expansion) is indeed a bona fide killed diffusion.

#### S2.1 Deriving the infinitesimal generator of the killed diffusion

We aim to find the generator of the stochastic process defined by Eq. S7. Let Ω := [0, 1] ∪ {*C*} be the state space of our stochastic process. Note that *P*( *f, t*) vanishes at the cemetery state (by definition). We begin by multiplying both sides of Eq. S7 by an arbitrary smooth function *ψ* with compact support^3^ in Ω and integrating over Ω:

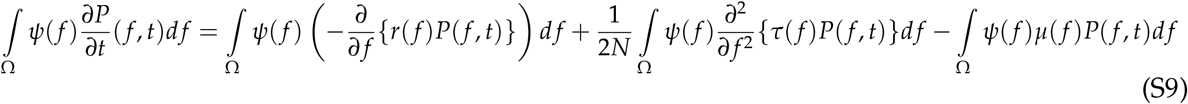

We now use integration by parts on the first two terms of the RHS and discard the boundary terms since *ψ* has compact support. Doing this once on the first term of the RHS and twice on the second term of the RHS yields

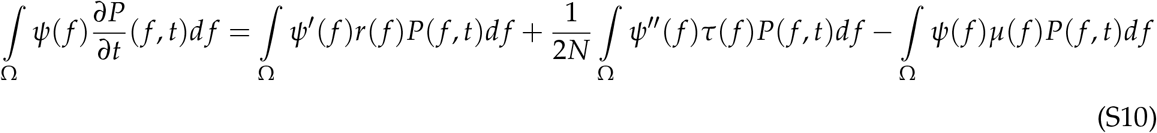

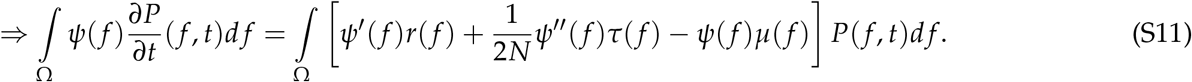

Now, by the definition of the derivative, the LHS of Eq. S11 is

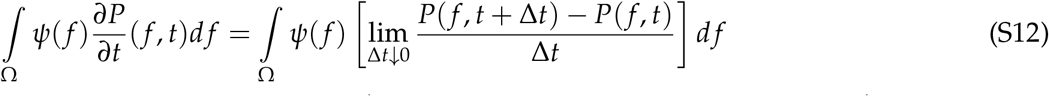

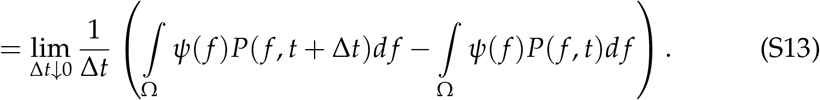

Interchanging the order of limits and integrals is permissible by the dominated convergence theorem. The integrals on the RHS of Eq. S13 are, by definition, expectation values. We have thus obtained

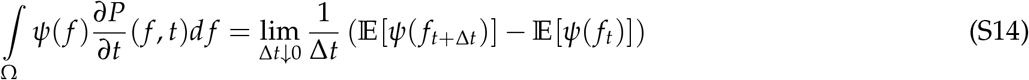

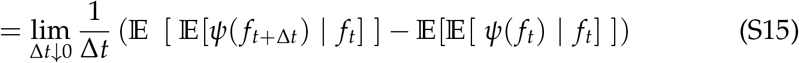

where we have used the tower property E[*X*] = E[ E[*X* |*Y*] ] to obtain the last equality. Using the linearity of the expectation value and once again interchanging the order of limits and integrals (expectations) gives us

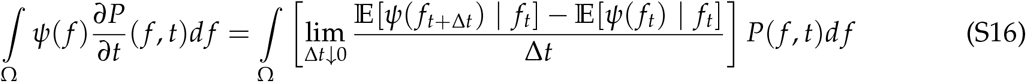

Replacing the LHS of Eq. S11 with the RHS of Eq. S16 now results in

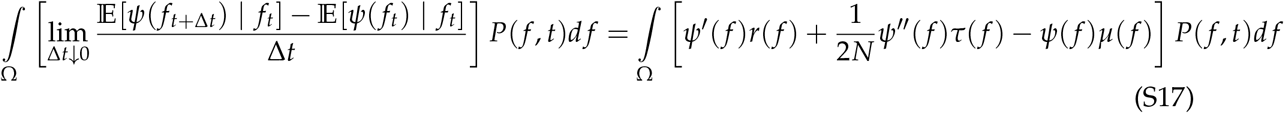

Since Eq. S17 is true for *every* (smooth, compactly supported) function *ψ*, we conclude that the generator *L* of our process is given by

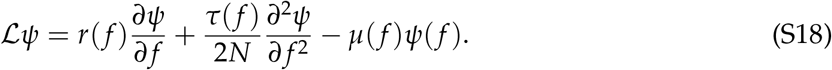

Besides being useful for further calculations, Eq. S18 also verifies that the process we have obtained after the diffusion approximation is indeed a killed diffusion (compare our Eq. S18 with Eq. 2 in Steinsaltz and Evans, 2006).

#### S2.2 A Feynman-Kac representation for functions of the killed diffusion

The infinitesimal generator of a Markov process is particularly useful to us due to a result called *Kolmogorov’s backward equation* (Øksendal, 1998, Theorem 8.1.1; Karlin and Taylor, 1981, Eq. 15.5.7).

Let {*X*_*t*_}_*t*_ be a time homogeneous Markov process on a nice domain *D* ⊆ ℝ. Let *G* denote the generator of *X*, and consider a real function *g* ∈ *C*^2^(*D*). Let *u*(*x, t*) := E[*g*(*X*_*t*_)|*X*_0_ = *x*] denote the expected value of *g*(*X*) after time *t* given that the process {*X*_*t*_}_*t*≥0_ began at state *x*. Kolmogorov’s backward equation (Øksendal, 1998, Theorem 8.1.1) says that *u*(*x, t*) satisfies

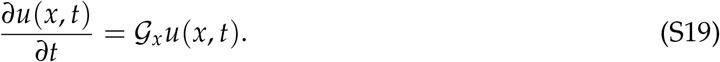

The notation *G*_*x*_ is to indicate that the generator *G* is acting on the map *x* ↦ *u*(*x, t*). Notice that while the LHS of Eq. S19 is about the state at time *t* (forward in time), the RHS of Eq. S19 is in terms of changes to the *initial condition x* (contrast with, say, Eq. S7, where derivatives are with respect to the future/current state).

For our killed diffusion, the generator is Eq. S18. Let 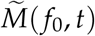 denote the expected value of a measurement 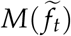 given an individual is born with *f*_0_ failures at age 0 and follows the killed diffusion 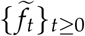 defined by Eq. S18. Here, we specify that we require^4^ *M* ∈ *C*^2^ ([0, 1]). Kolmogorov’s backward equation states that 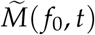 satisfies the initial value problem

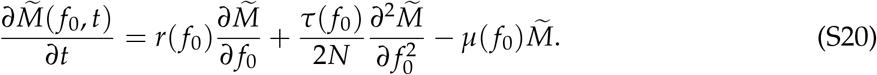

with the initial condition 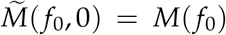. The Feynman-Kac formula in the PDE to SDE direction (Øksendal, 1998, theorem 8.2.1, (b)) now states that since 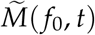 satisfies Eq. S20, it must admit the representation

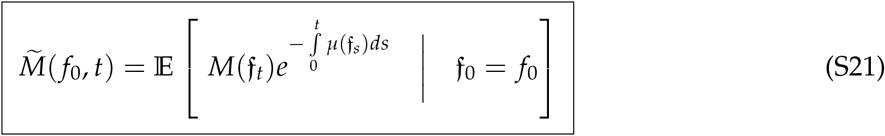

where {*f*_*t*_}_*t*≥0_ is the process described by the Tithonus model that solves the SDE Eq. 1. The conditional expectation in Eq. S21 is understood as an expectation over paths (*i.e*. on the space {*ω* ∈ *C*([0, *t*], [0, 1]) | *ω*(0) = *f*_0_} equipped with the Wiener measure induced by the diffusion f_*t*_). We also explain an intuitive way to numerically calculate such expectations in the next sub-section.

#### S2.3 Conditional expectations over the paths of the solution to an Itô SDE

Consider a continuous time stochastic process {*X*_*t*_}_*t*≥0_ that takes values in [*a, b*] ⊆ ℝ ∪ {−∞, ∞} and solves the SDE *dX*_*t*_ = *F*(*X*_*t*_)*dt* + *G*(*X*_*t*_)*dW*_*t*_. Let *H*(*X*_*T*_) be a function that depends on the entire path or ‘history’ of {*X*_*t*_}_*t*≥0_ up to the time *T* > 0. For instance, the survivorship function 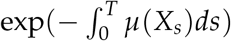 is one such function. One way of understanding the meaning of the conditional expectation E[*H*(*X*_*T*_) | *X*_0_ = *x*] is to partition [0, *T*] into *n* discrete time intervals, each of width (Δ*t*)_*n*_ := *T*/*n*. For each *k* ∈ {0, 1, …, *n*}, let *t*_*k*_ := *kT*/*n* denote the *k*^th^ timepoint in the partition. Note that *t*_0_ = 0 and *t*_*n*_ = *T*. The conditional expectation of *H*(*X*_*T*_) over the paths of the Itô diffusion *X*_*t*_ can be written as (Karatzas and Shreve, 1998, Theorem 2.4.20 with Theorem 3.5.1)

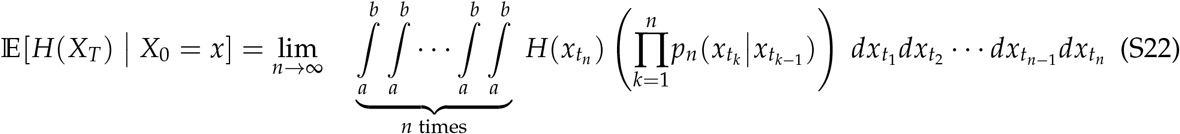

where we set 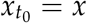 to enforce the conditioning on the initial value, and each

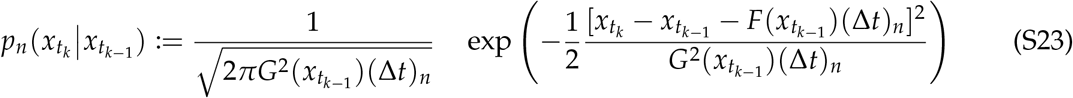

can be interpreted as the density function for the process {*X*_*t*_}_*t*≥0_ to take value 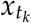 at time *t*_*k*_ given that its value at an earlier time *t*_*k*−1_ was 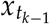. The product 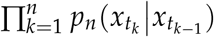 is thus just a joint density function for 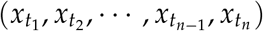, a vector that becomes a continuous path as *n* → ∞, *i.e*. as (Δ*t*)_*n*_ → 0. The fact that 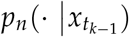 is the density function of a Normal distribution with mean 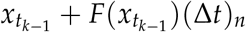 and variance 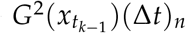 should make sense upon revisiting the meaning of the SDE *dX*_*t*_ = *F*(*X*_*t*_)*dt* + *G*(*X*_*t*_)*dW*_*t*_. The limiting object on the RHS of Eq. S22 can also be recast into a single, more abstract integral called an ‘integral with respect to the Wiener measure’ (Kac, 1949) in mathematics and a ‘Euclidean path integral’ (Feynman, 1948) in physics.

Eq S22 also provides a method to numerically calculate conditional expectations such as those that occur in Eq. S21 (and Eq. 6 in the main text). To calculate the expectation, we only need to numerically the densities Eq. S23, which is easily done by directly simulating a large number of Tithonus paths via standard SDE simulation tools such as the Euler-Maruyama method (see e.g. Gardiner (2009), Chapter 15).

### S3 Feynman-Kac representation of the killed diffusion conditioned on remaining alive

Yashin et al., 1985 have shown that if we know that the processs enters the cemetery state at time *T*_mort_, the conditional density 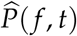 of the process conditioned on not entering the cemetery state (*t* < *T*_mort_) obeys the Fokker-Planck type PDE

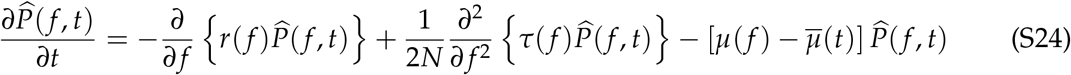

where

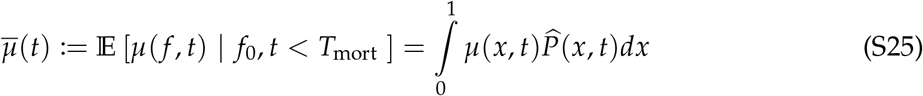

is the expected force of mortality conditioned on remaining alive. Feynman-Kac formulae cannot be used directly for the conditioned process defined by Eq. S24 because Eq. S25 introduces a non-linearity in the probability density 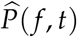. In this section, we use Eq. S21 to derive a representation instead.

From Eq. S21, we found that the survival function 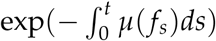 provided the correct weighting to modify expectations from the Tithonus model to find expectations of the process with disappearance. Since conditioning effectively modifies the mortality term *µ*( *f* ) → *µ*( *f* ) − *µ*(*t*) in the Fokker-Planck equations (compare Eq. S24 with Eq. S7), we may guess that the survival function for the conditioned process should also be modified as 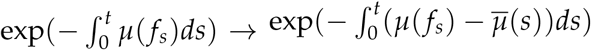. To this end, let^5^ 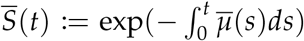, and consider 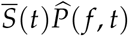. Differentiating with respect to time yields

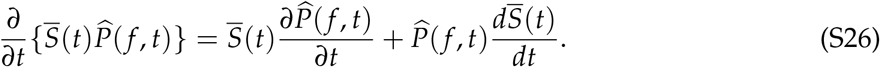

Now,

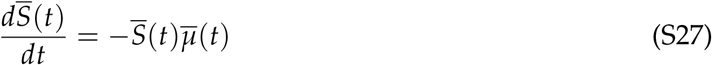

and 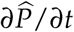 is given by Eq. S24. Substituting into Eq. S26, the 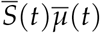 terms cancel (since 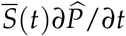 contains a 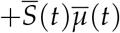 term) and we find

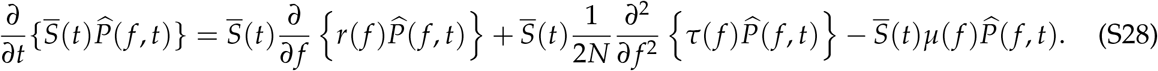

Since 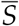 is independent of *f*, it can be taken inside the partial derivatives on the RHS, resulting in

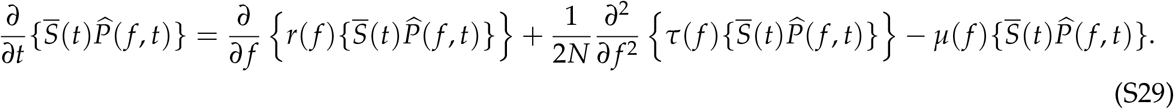

Comparing Eq. S29 with Eq. S7, we find that 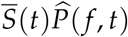 is precisely the probability density function of the *unconditioned process* with disappearance! Letting 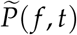 denote the density of the unconditioned process (for consistency with the main text), we have thus shown that

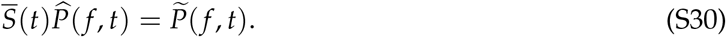

Eq. S30 is a powerful result because it establishes a relation between the unconditioned and conditioned processes at the level of their density functions. It also establishes that 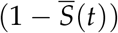 quantifies the loss of density to the cemetery state in the unconditioned process. However, 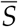 still involves the conditioned density 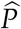 via the non-linear term Eq. 5. We seek a better representation of 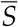 in terms of the Tithonus model.

Begin by integrating both sides of Eq. S30 over all possible values of *f* . Since 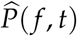 is a probability density function for a process that is conservative in the failure space, it integrates to 1 (intuitively because we conditioned on not dying and so the individual must have some value of *f* ; also provable directly from Eq. S24, see Yashin et al., 1985). This results in the expression

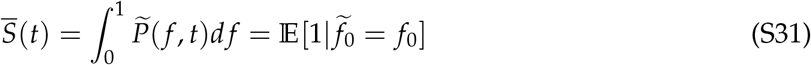

where 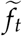 is a stochastic process with generator Eq. S18 (*i.e*. following the unconditioned process with disappearance) and the expectation on the RHS is not equal to 1 because individuals are lost to the cemetery state (*i.e*. the process is non-conservative).

We now want to represent the RHS of Eq. S31 in terms of the process described by the Tithonus model. To do this, we notice that the RHS of Eq. S31 is a special case of 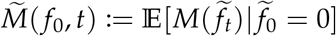 with the choice of measurement function *M*(·) = 1. Thus, from Eq. S21 with *M* ≡ 1, we find

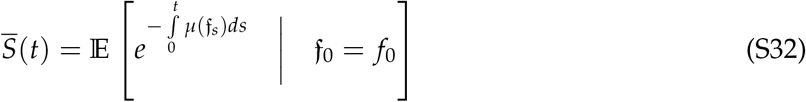

where {f_*t*_}_*t*_, like in Eq. S21, solves the SDE Eq. 1.

To find 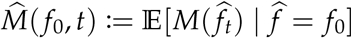, the expected number of failures accrued by an individual conditioned on survival (*i.e*. following the conditioned process defined by Eq. S24), multiply both sides of Eq. S30 by *M*( *f* ) and then integrate over all possible values of *f* . This yields

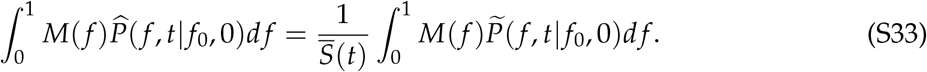

But by definition, the integral on the LHS is 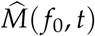 and the one on the RHS is 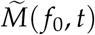. Substituting Eq. S21 and Eq. S32 now finally yields

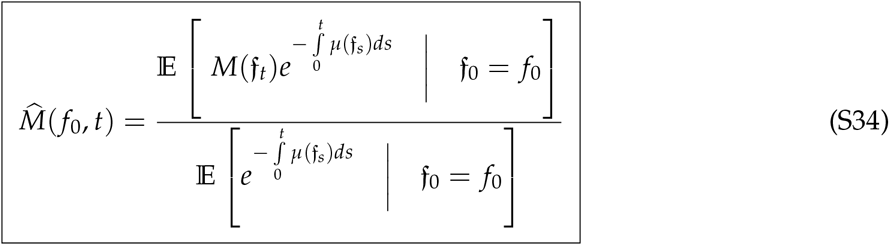

The RHS of Eq. S34 is entirely in terms of the Tithonus SDE f_*t*_ and is Eq. 6 in the main text.

### S4 Bounding the expected deviation in a measurement due to selective disappearance

We begin by squaring both sides of Eq. 8, thus finding that the squared discrepancy is given by

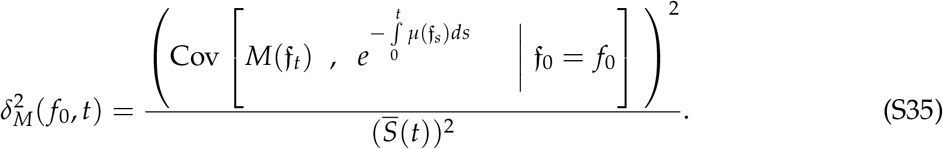

For convenience, we henceforth denote conditional expectations, variances, and covariances by E _*f*_, Var _*f*_, and Cov _*f*_ respectively. Let^6^ 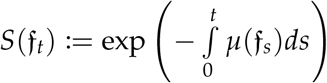 . The Cauchy-Schwartz inequality immediately lets us bound the numerator of the RHS of Eq. S35,

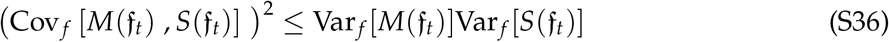

leading to

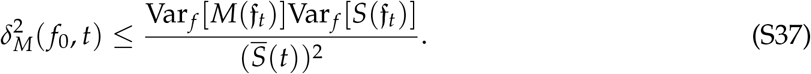

We now bound the second variance in the numerator of the RHS of Eq. S37. By definition,

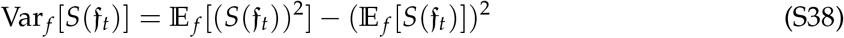

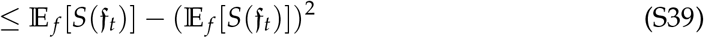

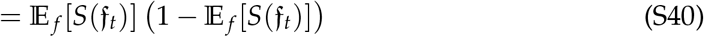

where we have used the fact that 0 ≤ *S*(f_*t*_) ≤ 1 and thus *S*^2^ ≤ *S* to go from Eq. S38 to InEq. S39. Now notice that E _*f*_ [*S*(f_*t*_)] = *S*(*t*) from Eq. S32, and we hence have

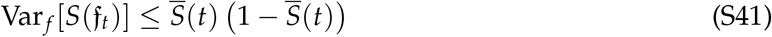

From InEq. S41 and Ineq. S37, we can conclude that

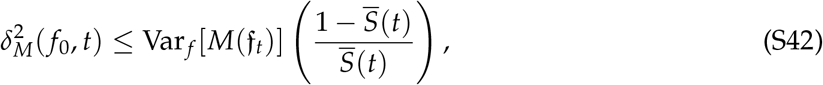

*i,e*.,

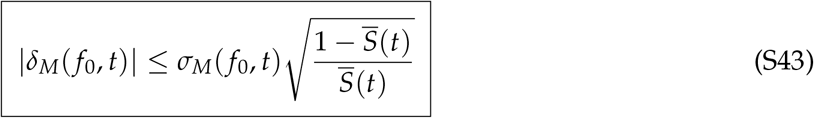

where 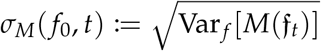 is the standard deviation process of {*M*(f_*t*_)}_*t*≥0_. This is Ineq. 9 in the main text.

We will now further bound the standard deviation on the RHS of Eq. S43 for the special case *M*(·) = *µ*(·). In this case, we know that *µ*(f_*t*_) solves the SDE Eq. 2, allowing us to write

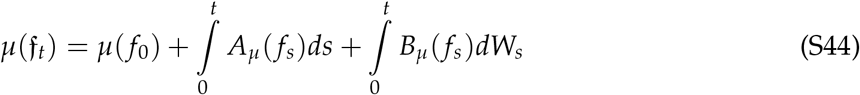

where 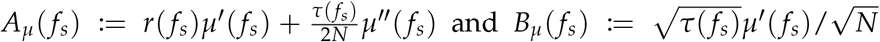 The Itô isome-try (Karatzas and Shreve, 1998, Chapter 2, Proposition 2.10) then gives us

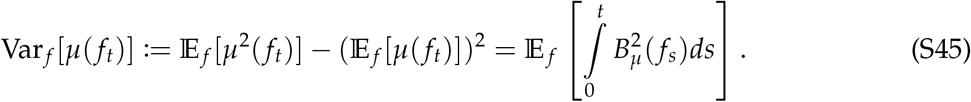

We now assume that both *τ*( *f* ) and *µ*^′^( *f* ) are bounded above. In other words, we assume that there are finite numbers 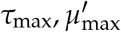 such that *τ*( *f* ) ≤ *τ*_max_ and *µ*^′^( *f* ) ≤ *µ*_max_. In this case, we have

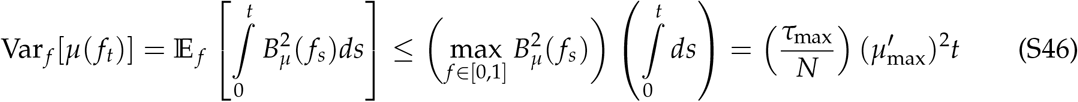

and can hence conclude that

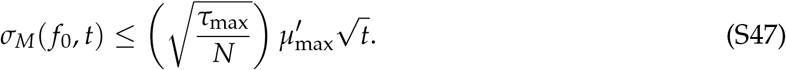

We can now plug Ineq. S47 into Ineq. S43 to arrive at

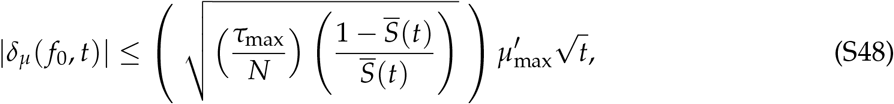

which is Eq. 10 in the main text. As an aside, note that the same idea would carry through to bound |*δ*_*M*_| for any *C*^2^([0, 1]) measurement function *M* thanks to Itô’s formula as long as the absolute value of the derivative of *M* is bounded above^7^ by 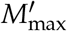. The only change will be that 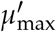 will be replaced by 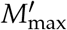 in Eq. S48. If we were interested in, say, fecundity, for example, the relevant bound would be the maximum rate at which fecundity changes as a function of failures (assuming that fecundity as a function of failures is a twice continuously differentiable function).

### S5 Bounding the expected deviation from Tithonus mortality curves when the geometric mortality rule is obeyed

The geometric mortality rule Eq. 13 that we advocated for in the main text does *not* have a bounded derivative, since our model predicts infinitely high mortality risk when *f* = 1. Happily for us, 0 ≤ *f* ≤ 1, and we can thus use a geometric series expansion for *f* /(1 − *f* ). In other words, our mortality rule can be rewritten as

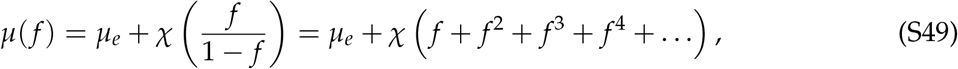

which will be bounded for any finite truncation of the infinite sum. Truncating Eq. S49 at a finite degree *α* > 1 results in the approximate mortality rule

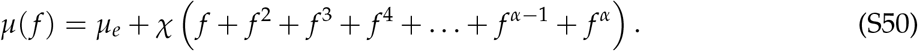

Higher values of *α* will provide strictly better approximations to the mortality rule Eq. 13 used in the main text, and the equality is exact if we let *α* → ∞. For finite *α*, we can calculate and bound the derivative of Eq. S50, since

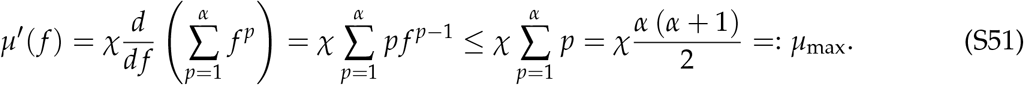

Plugging into Ineq. S48 now reveals that

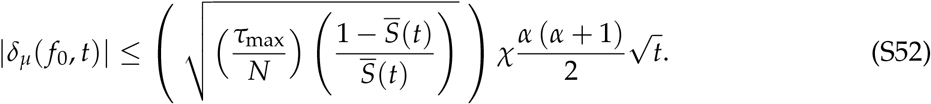

Finally, we will make the dependence on extrinsic mortality explicit. Notice that

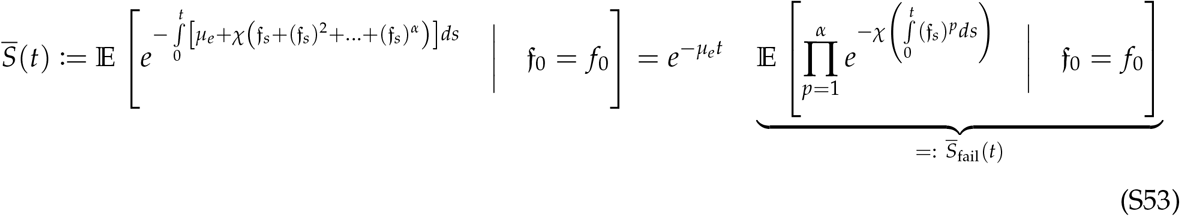

where 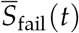 quantifies the survivorship when ignoring all deaths due to extrinsic mortality (we can do this because we have constructed the mortality rule such that it is the sum of an extrinsic and an intrinsic component). Substituting into Ineq. S52 now yields:

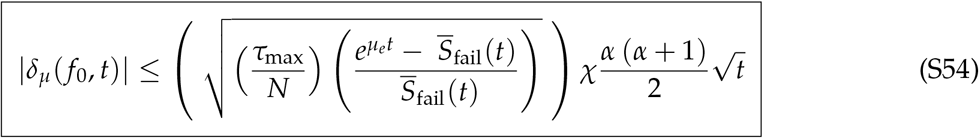

where 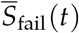 measures the expected survivorship when deaths due to extrinsic factors are excluded. In other words, we have partitioned the overall expected survivorship 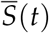 from Eq. 7 as 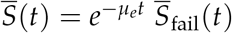. Ineq. S54 tells us that

1. Deviations caused by selective disappearance are less important at smaller ages, but can become more important at later ages. This simply reflects the fact that the role of heterogeneity and ‘luck’ in affecting distributions via disappearance is more important as the cohort ages.
2. Selective disappearance becomes less powerful at causing deviations from Gompertz as the total number of sub-systems *N* grows larger. As mentioned in the main text, the effect of increasing *N* can be thought of as increasing the ‘total number of ways you can die of sub-system failure’.
3. Selective disappearance becomes less important as *τ*_max_ becomes smaller. Recalling that *τ*( *f* ) controls the variance in the failure accumulation process over small timescales, this means that selective disappearance becomes less important as the failure accumulation process itself becomes more stereotyped/canalised.
4. For a fixed value of *µ*_*e*_, the role of extrinsic mortality in causing deviations due to selective disappearance is more important at later ages. Conversely, for a fixed age, selective disappearance becomes less important as the extrinsic mortality *µ*_*e*_ decreases. Though this may seem puzzling at first glance because extrinsic mortality traditionally masks patterns of senescence, it becomes more clear once we remember that we are calculating conditional probabilities. Intuitively, extrinsic mortality uniformly controls the probability of survival up to a certain age, independent of failures. Since we are conditioning on individuals remaining alive up to age *t*, this conditioning increasingly focuses on those ‘lucky’ individuals who avoided death as (a) extrinsic mortality increases, and (b) later ages are considered. In other words, though extrinsic mortality does not itself cause ‘selective’ disappearance in the sense of favouring some states over others, its effects are nevertheless to cause deviations from curves predicted without disappearance because extrinsic mortality is selecting for pure dumb luck. However, this is only true when failures do still have an impact on mortality (note the *χ* outside the square root) because extrinsic mortality can only amplify existing differences.
5. Selective disappearance becomes less visible as 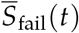, the expected survivorship when deaths due to extrinsic causes are ignored, increases. The same logic also explains the multiplicative factor of *χ* outside the square root: if failures have a smaller impact on mortality overall (*χ* is smaller), selective disappearance is less visible; if all mortality is extrinsic and failures do not impact mortality at all (*χ* = 0), selective disappearance cannot cause any deviations of mortality curves from the predictions of the Tithonus model.

### S6 A mean field failure accumulation rule when sub-systems are vertices of a vertex-transitive graph

In this section, we sketch how a plausible failure accumulation rule can be derived for organisms in which the sub-systems are organised in an (extremely symmetric) graph/network. This model is inspired by Nielsen et al. (2024).

#### S6.1 Definition of the process

We conceptualize an organism as a simple undirected connected graph *G* on *N* vertices. Each vertex of G is a sub-system important for organismal function, and edges represent interdependencies between systems. We assume that the graph *G* is *vertex transitive*, meaning that the vertices of the graph are indistinguishable (Godsil and Royle, 2001, section 3.1; For an application of such graphs in evolutionary biology, see McAvoy and Hauert, 2015). Biologically, this assumption is a symmetry assumption where we are assuming that every sub-system is equally important and structurally indistinguishable from the other. Since every vertex transitive graph is regular (Godsil and Royle, 2001, Section 3.1), our assumption also implies that every sub-system (vertex) has the same number of dependencies (edges), a quantity we denote by *n*. While vertex transitivity is a strong assumption, it is not overly restrictive for a first pass, since the number of connected vertex-transitive graphs on *N* vertices grows very quickly with *N*. For instance, there are 34333611 connected vertex transitive graphs on *N* = 47 vertices (OEIS A006800).

Failure accumulation is modelled as a continuous-time stochastic process that consists of relabelling the vertices of *G* as failed/non-functional. In particular, if a vertex *v* is functional, we assume it has a failure rate *ϕ* + *In*_*v*_(*t*), where *ϕ* > 0 is a constant intrinsic failure rate due to factors like physical damage and wear and tear and *n*_*v*_(*t*) is the number of neighbours of *v* that have failed at time *t*. Here, *I* > 0 is a parameter that controls how much a focal subsystem is affected by the failure of the subsystems on which it depends. The special case *I* = 1 corresponds to the case where failure depends on the number of failed neighbours. Let ℙ(*E*) denote the probability of the event *E* occurring. For conciseness, let us use the shorthand *f* (*i*; *j*) for the rate at which the *i* th vertex fails when it has *j* failed neighbours, i.e.

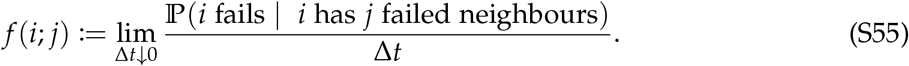

Our model above specifies the particular rule *f* (*i*; *j*) = *φ* + *I j*, but we change notation here because we will be summing over *i* and *j*. Letting *F*_*t*_ be the stochastic process tracking the number of failures by time (age) *t*, we are interested in computing the failure rate

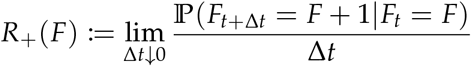

with the ultimate goal of relating it to the s defined in the main text. Repair rates can be computed exactly analogously and we thus do not repeat the calculations.

#### S6.2 Computing the failure rate

By definition, the number of failed systems *F*_*t*_ increases iff a functional system fails. Thus, the rate at which the number of failed sub-systems *F*_*t*_ increases is given by the rate at which at least one functional vertex fails. In equations, we express this as

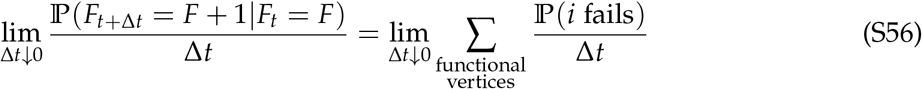

where the sum is over the set of sub-systems that have not yet failed, and we have neglected the possibility of two vertices failing at the same time because this is an *O*((Δ*t*)^2^) event. Let Δ_*i*_ denote the (in)-degree of vertex *i* (*i.e*. the number of systems that *i* depends on). We can rewrite the probability on the RHS of (S56) as

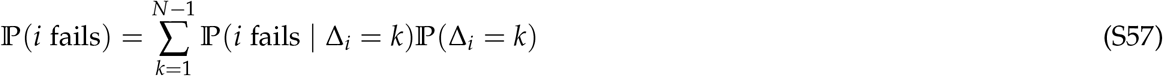

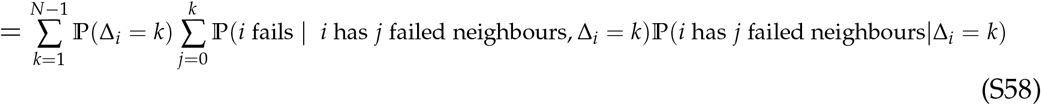

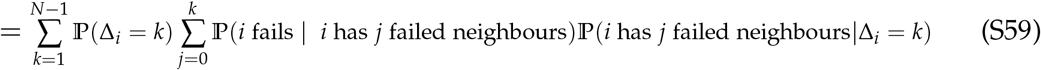

Thus, we have

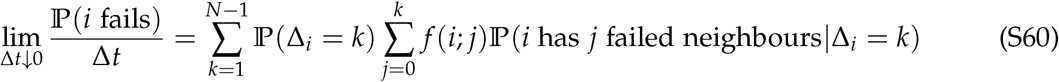

where we have used the definition of *f* (*i*; *j*). We now observe that ℙ (*i* has *j* failed neighbours|Δ_*i*_ = *k*) is the probability that a vertex has *j* failed neighbours given that it has *k* neighbours. Assuming (in the mean-field) that the probability that a given randomly chosen vertex is a failed vertex is *F*/*N* (the fraction of failed vertices), ℙ(*i* has *j* failed neighbours|Δ_*i*_ = *k*) must follow a Binomial(*k, F*/*N*) distribution. Thus, we can rewrite Eq. (S60) as

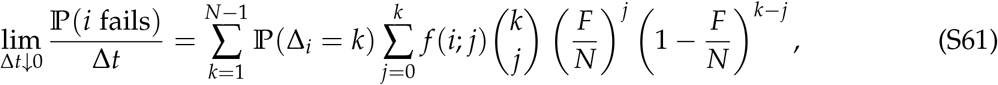

which, upon substituting into Eq. (S56) yields the failure rate

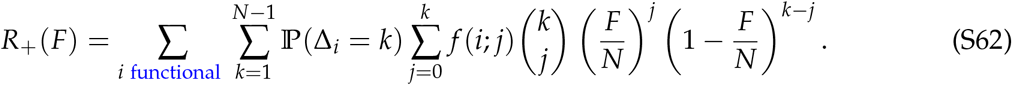

If vertices are indistinguishable, *i.e*. the process on every vertex *i* is identical, Eq. (S62) further simplifies to

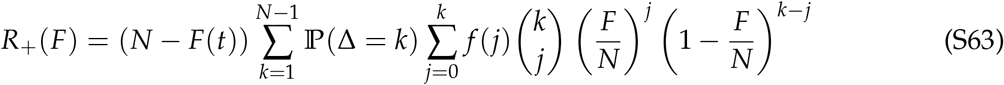

We will now substitute the functional form *f* (*i*; *j*) = *φ* + *I j*. Upon doing this, Eq. (S63) becomes

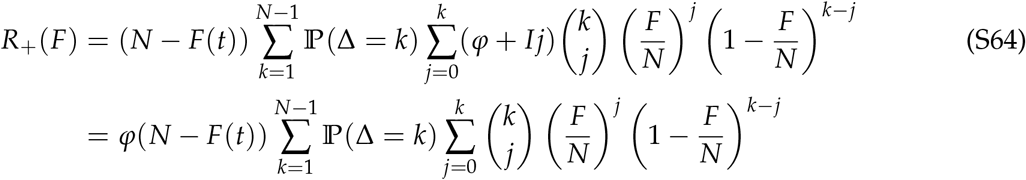

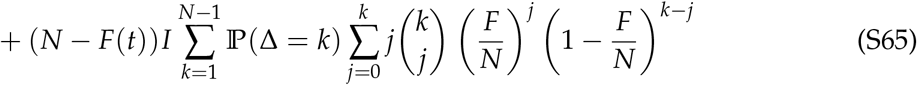

We now observe that both sums over *j* are expressions for moments of the binomial distribution. If *X* is a Bin(*n, p*) random variable, we have

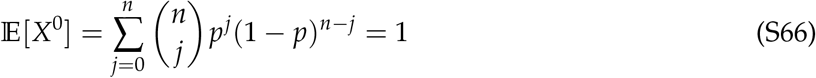

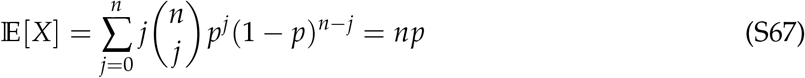

In our case, we have *n* = *k* and *p* = *F*/*N*. Using this observation in Eq. (S65), we obtain

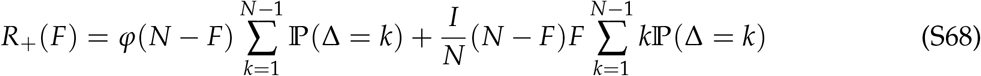

The two sums on the RHS of Eq. (S68) are now just moments of the degree distribution. The first sum simply equals 1, and the second is the average degree ⟨Δ⟩. We are thus led to the failure

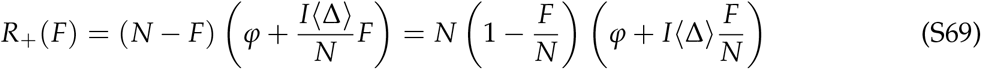

Defining *f* = *F*/*N*, we see that our graph-structured model corresponds to the rule *r*_+_( *f* ) = (1 − *f* )(*φ* + *I*⟨Δ⟩ *f* ). Since the ability to repair failed sub-systems *decreases* with the proportion of failed sub-systems, the logic for deriving the repair rate *r*_−_( *f* ) is exactly analogous, and will just have different constants *φ*_2_ and *I*_2_. Thus omitting the derivation for the repair rate *r*_−_( *f* ), we can identify that our per-capita failure accumulation rate is *ρ*( *f* ) = *φ*_eff_ + *I*_eff_⟨Δ⟩ *f*, leading us to conclude that *ϕ*_*ρ*_ = *φ*_eff_, *k*_*ρ*_ = *I*_eff_⟨Δ⟩. Thus, interdependence *k*_*ρ*_ is a constant times the average degree ⟨Δ⟩, an explicit measure of the number of sub-systems the average sub-system depends on.

**Remark**

If the failure rate of a vertex depends on the *proportion* of failed neighbours rather than the number of failed neighbours, we would have

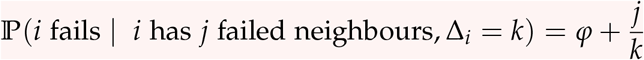

in Eq. S58. Using this new relation, the last term on the RHS of Eq. S65 is instead

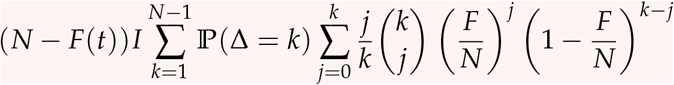

and the last term on the RHS of Eq. S68 hence becomes

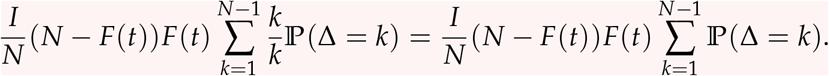

Thus, the rate *R*_+_(*F*) in this case becomes

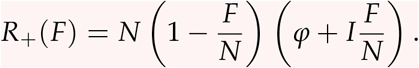

In other words, unlike in Eq. S69, the network structure (parametrised by the average degree ⟨Δ⟩ = *n*) does not affect the mean field dynamics if failure rate depends on the proportion of failed neighbours.

### S7 Given a mechanistic failure accumulation rule, what sorts of mortality rules generate Gompertz-Makeham curves in the Tithonus model?

In the previous section, we formulated a mechanistic model of failure accumulation in which an organism is viewed as a (very symmetric) graph, with vertices representing sub-systems and edges representing interdependencies. We then showed that this model leads to the null model we called ‘logistic failure accumulation’ in the main text. In the main text, we also argued that a general class of failure accumulation models should look approximately like the logistic failure accumulation rule during the initial part of the failure accumulation process. This last argument essentially arises from a linearisation of the per-capita failure accumulation rate *ρ*(*f* ). We then showed (in the main text) that the geometric mortality rule combines with the logistic failure accumulation rule to generate Gompertz-Makeham mortality curves. In this section, we ask a slightly different question: If you are given a mechanistic (possibly quite complex) failure accumulation rule based on some known underlying biological principles, what must the mortality rule *µ* look like to ensure that the emergent mortality curve (in the Tithonus setting with infinitely many sub-systems) is Gompertz-Makeham?

We assume the failure accumulation curve is the solution to the ODE

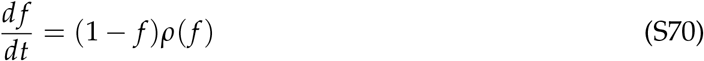

In practice, *ρ*( *f* ) should be derived from a biological basis via a mechanistic model such as the network-structured model of section S6 or other detailed reliability-theoretic models (see e.g. Gavrilov and Gavrilova, 1991), and thus may be quite complicated with many high-order non-linearities. We assume that *ρ* is smooth, *ρ*( *f* ) > 0, *ρ*^′^( *f* ) > 0, and that *ρ*(1) and *ρ*^′^(1) are finite.

We parameterise the mortality rule as

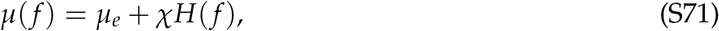

where *µ*_*e*_, *χ* > 0 are constants and *H*( *f* ) is a function (to be found) describing the additional mortality hazard due to failures. The geometric mortality rule, in this view, is the specific choice *H*( *f* ) = *f* /(1 − *f* ).

#### S7.1 Gompertz-Makeham implies an inverse power-law scaling of the mortality rule (the ‘only if’ direction)

Assume that the Gompertz-Makeham law is obeyed, i.e. that the overall mortality curve *µ*(*t*) as a function of age from the Tithonus model is known to be described exactly by

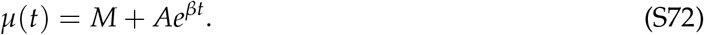

for known or estimated parameters *M, A*, and *β*. We are interested in determining what *H* must necessarily look like given these facts. Equating the RHS of Eq. S71 with the RHS of Eq. S72, we find

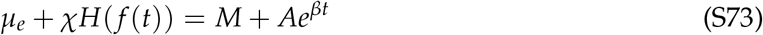

where *f* (*t*) solves the ODE Eq. S70.

Now, since we assume throughout that *ρ*( *f* ) > 0 ∀ *f* ∈ [0, 1), Eq. S70 leads to *f* monotonically increasing with age. Consequently, we can invert the expression for *f* (*t*) to find the age *T*( *f* ) at which an organism accumulates *f* failures. Doing this from Eq. S70 gives

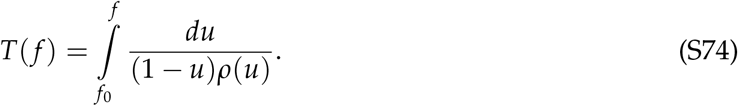

Since *f* (*t*) in Eq. S73 obeys Eq. S70, *t* in the RHS of S73 must be given by the RHS of Eq. S74 (i.e. *t* = *T*( *f* (*t*))). Thus, we find (after some rearrangement) that we must have

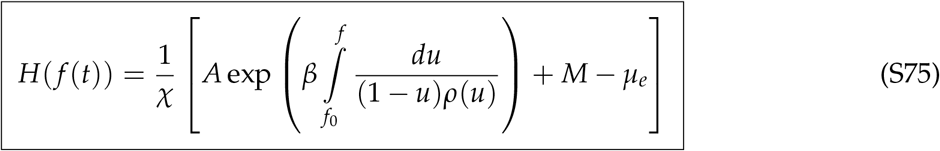

If both (i) a precise, mechanistically motivated failure accumulation rule *ρ*, and (ii) estimated Gompertz-Makeham parameters *M, A, β* are provided on empirical grounds, Eq. S75 (with Eq. S71) makes an exact theoretical prediction for the connection between failures and mortality, since the *H* that satisfies Eq. S75 is unique^8^.

Though Eq. S75 precisely characterises the required mortality rule, it is not a very transparent expression. To better understand the behaviour of the mortality hazard *H*, we need to understand the integral

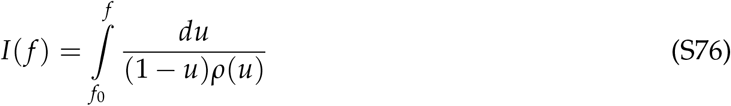

We will develop a picture of the functional form of *H* by examining the behaviour of this integral at the extreme value *f* = 1 (close to complete sub-system failure). Since we are interested in asymptotics and functional forms rather than precise numerical values, we will use *C*_*i*_ to denote constants whose precise value is not of interest to us and may change from line to line.

##### S7.1.1 The behaviour close to complete failure

The behaviour when *f* is close to 1 is trickier to study because the integrand has a singularity at *f* = 1, but is also more insightful about the allowed functional forms of *H*. The (asymptotic) failure accumulation rate at *f* = 1 is given by *ρ*(1). Biologically, *ρ* is the per-capita failure accumulation rate of a physiological sub-system, and we expect it to always remain finite and non-zero in (0, 1). We assume that the limiting failure accumulation rate *ρ*(1) is also finite (and non-zero). We wish to study the asymptotic behaviour of *I*( *f* ) very close to *f* = 1, accounting for the singularity due to the 1/(1 − *u*) term in the integrand. We will do this by “isolating” the point singularity at 1. To this end, pick a sufficiently small *ϵ* > 0, and let *f* ∈ (1 − *ϵ*, 1). We can split the required integral into two terms

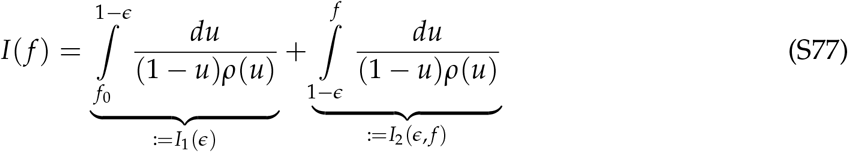

where *I*_1_ depends only on *ϵ* and is guaranteed to be finite. Now, since *ρ* is continuous, we can approximate *ρ*( *f* ) by *ρ*(1) in the region (1 − *ϵ*, 1) for sufficiently small *ϵ* (we will soon be taking the *ϵ* → 0 limit). Thus, the second integral can be evaluated as

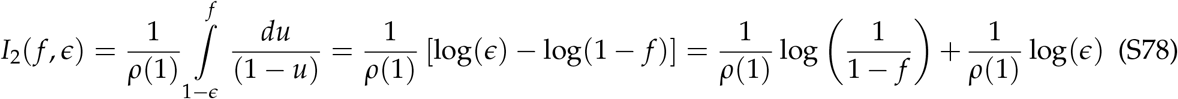

To evaluate the first integral, we use a method of partial fractions. Anticipating that the *ϵ* dependence of *I*_1_ should cancel out with the *ϵ* dependence of *I*_2_ in Eq. S78 (since the LHS of Eq. S77 is independent of *ϵ*), we add and subtract 1/(*ρ*(1)(1 − *u*)) within the integrand of *I*_1_. This yields

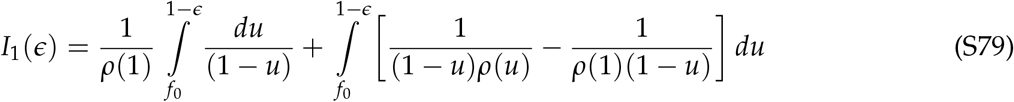

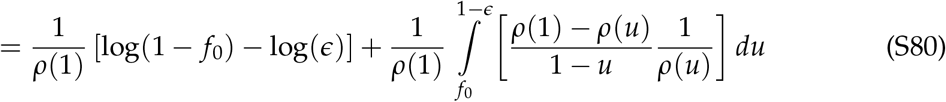

As expected, the − log *ϵ* term in Eq. S80 will perfectly cancel with the + log *ϵ* term in Eq. S78 when *I*_1_ and *I*_2_ are added to find our focal integral.

Thus, asymptotically close to *f* = 1, (i.e. for *f* ∈ (1 − *ϵ*, 1] as *ϵ* → 0), we find

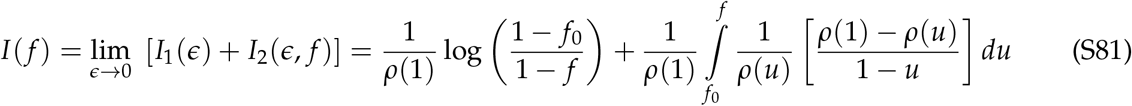

where we have used the fact that since *f* ∈ (1 − *ϵ*, 1], the limit of integration 1 − *ϵ* in the rightmost integral of Eq. S80 can be replaced by *f* as *ϵ* → 0. Now observe that since 0 < *ρ*(*u*) < ∞ and the rightmost integrand of Eq. S81 is bounded in [ *f*_0_, 1] as long as^9^ *ρ*^′^(1) < ∞, it just converges to some finite constant,

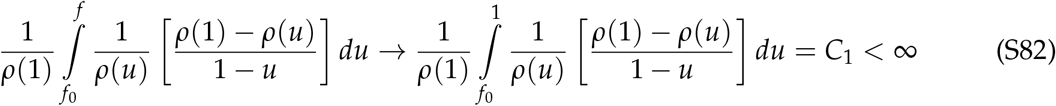

Very close to *f* = 1, our focal integral *I*( *f* ) must thus behave like

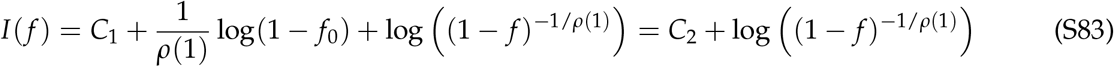

Substituting into the equation of our failure-dependent mortality hazard Eq. S75, we have thus found that *H*( *f* ) must asymptotically show the behaviour

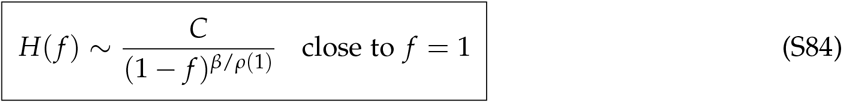

Thus, if the emergent mortality curve for a model with per capita failure accumulation function *ρ* is known to be Gompertz-Makeham with rate parameter *β*, the failure-dependent part of the mortality rule must show inverse power-law like behaviour near *f* = 1, with a singularity at *f* = 1 and the power-law scaling exponent *γ* := *β*/*ρ*(1).

#### S7.2 Inverse power law scaling of the mortality rule implies asymptotically Gompertz mortality (the ‘if’ direction)

If a mortality rule obeys Eq. S75 for *some A, β* > 0, the Gompertz-Makeham law Eq. S72 will of course be exactly obeyed because we can simply make the calculations from Eq. S75 to Eq. S73 in reverse order. However, Eq. S75 is quite restrictive and it is not clear why it should hold in general. In this section, we will prove a weaker asymptotic result for a larger class of mortality rule using what we learnt about the necessary conditions for the Gompertz-Makeham law to be obeyed.

Suppose we have the failure accumulation rule Eq. S70 and a mortality rule Eq. S71. Assume that *H*( *f* ) ∼ *C*_3_(1 − *f* )^−*γ*^ as *f* → 1^−^, for a constant *C*_3_ > 0 and a scaling exponent *γ* > 0. We will show that given sufficiently advanced ages, the mortality curve with age, *µ*(*t*), will obey the Gompertz law.

Begin with the failure accumulation dynamics Eq. S70, and consider the variable *x* = 1 – *f* tracking the fraction of healthy sub-systems. From Eq. S70, *x* obeys the ODE

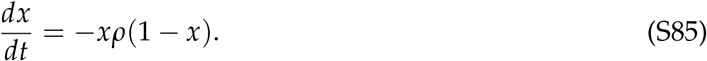

Now, by assumption, following an initial (possibly long) transient, *f* → 1 (because *ρ*( *f* ) > 0 ∀ *f* ) and thus *x* → 0. This means that asymptotically, given enough time, Eq. S85 relaxes to 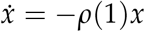, and thus, after a sufficiently large time *t*, we have

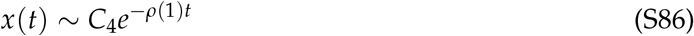

Now, by assumption, our mortality hazard *H*( *f* ) ∼ *C*(1 − *f* )^−*γ*^, which can be rewritten in terms of *x* as *H*(*x*) ∼ *Cx*^−*γ*^. Thus, given sufficient time, we have (using Eq. S86)

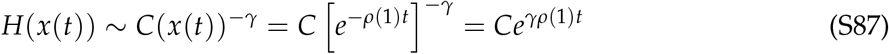

Using this in Eq. S71, we thus see that for sufficiently late ages, *µ*(*t*) ∼ *C* exp(*γρ*(1)*t*), which is the Gompertz law^10^ with ageing rate *γρ*(1).

Thus, if a model has a per-capita failure accumulation rate *ρ*( *f* ) and a mortality rule that asymptotically resembles (1 − *f* )^−*γ*^ for some *γ* > 0 as *f* → 1^−^, and the emergent mortality curve of the *N* → ∞ Tithonus model as a function of age will asymptotically become a Gompertz curve with rate parameter *γρ*(1).

Importantly, we are explicitly *not* claiming that Gompertz or Gompertz-Makeham mortality will be observed at *all* ages if a mortality rule scales as (1 − *f* )^−*γ*^. Indeed, exact adherence to Gompertz-Makeham mortality at all ages requires *H* to be given exactly by Eq. S75. What we have proved is instead that any (monotonic, smooth) mortality rule that looks like *µ*( *f* ) ∼ *C*(1 − *f* )^−*γ*^ as *f* → 1^−^ will generate a mortality curve that *asymptotically* relaxes to a Gompertz curve given sufficiently advanced ages, possibly after a non-exponential transient at earlier ages. An instructive example of the difference between asymptotically Gompertz and exact Gompertz-Makeham curves is the combination of the logistic failure accumulation rule *ρ*( *f* ) = *ϕ* + *k f* with the mortality rule that satisfies *H*( *f* ) = *f* /(1 − *f* )^2^. This system has *ρ*(1) = *ϕ* + *k* and the scaling exponent *γ* = 2. The result is a mortality curve that initially rises faster than exponentially, but eventually relaxes to an exponential curve with rate 2*ρ*(1) at sufficiently large ages.

Notice that as long as the failure rule is increasing (*ρ*^′^ > 0) and satisfies 0 < *ρ*(1) < ∞, the mortality curve will asymptotically be Gompertz regardless of the details of the failure accumulation process. Thus, for instance, any increasing failure accumulation rule will, when combined with our geometric mortality rule, predict a mortality curve that eventually becomes a Gompertz curve. However, the *rate* of the asymptotic Gompertz depends on *ρ*(1), and is thus affected by the details of failure accumulation near complete failure ( *f* = 1). For instance, if *ρ* has a power series expansion 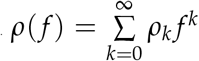, then 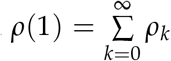. The logistic failure accumulation rule from the main text has *ρ*(1) = *ϕ*_*ρ*_ + *k*_*ρ*_, and the geometric mortality rule has *γ* = 1.

Another important caveat to remember is that all calculations of this section have been in terms of the *N* → ∞ Tithonus model, and the main model will still generate late-life plateaus via selective disappearance at later ages. Thus, determining whether the asymptotic convergence to a Gompertz is important requires information about the expected speed of convergence or the expected length of the non-Gompertzian transient. We suspect that Eq. S74, which expresses the age needed to reach *f* failures, could be used to estimate the expected duration of the non-Gompertzian transient for a given *ρ*( *f* ) and *µ*( *f* ). However, testing this suspicion and estimating the speed of convergence is beyond the scope of the present manuscript.

#### S7.3 The role of interdependence

Since the total rate of ageing is *γρ*(1), a higher value of *ρ*(1) leads to a faster rate of senescence if *γ* is held fixed. To make the connection with ‘interdependence’ *ρ*^′^( *f* ) of sub-systems as in the main text, observe that the fundamental theorem of calculus allows us to rewrite *ρ*(1) as

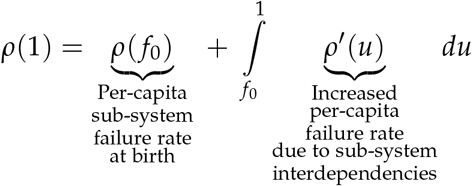

and, furthermore, we have argued that we must have *ρ*^′^ > 0 because failure begets failure. Thus, all else being equal, a uniformly greater interdependence *ρ*^′^( *f* ) will lead to a uniformly faster rate of senescence.

### S8 Explicit relation of our work to some previous mathematical models

In this section, we provide some explicit connections between our paper and some previous models of demographic senescence.

Our Eq. S7 is closely related to Woodbury and Manton’s (1977) ‘state space random walk’ model and extensions thereof (Woodbury and Manton, 1977, 1983; Manton et al., 1988; Mulder, 1993). In particular, by deriving diffusion equation descriptions from the ‘microscopic’ rules of failure accumulation and repair, our framework bridges mechanistic reliability-theoretic notions of senescence as damage/failure accumulation with the Woodbury-Manton view of senescence as a directed random walk/diffusion. Since Yashin et al., 1985 make the extension that is cleanest and most relevant to our work, we focus on connections with Yashin et al. (1985) here.

The exact correspondence between our work and that of Yashin et al., 1985 can be made by identifying their *a*(*t, y*) with our *r*( *f* ) and their *b*^2^(*t, y*) with our *τ*( *f* )/*N*. Unlike Yashin et al., 1985, however, we derive these coefficients and Eq. S7 more generally, from first principles starting from failure and repair of sub-systems (Eq. 3) as specified by a given mechanistic model (e.g. reliability theory). As such, we believe our derivation biologically grounds the coefficients that appear in Yashin et al. (1985) (their Eq. 5, our Eq. S7) in terms of failures of intra-organismal sub-systems. We also provide some guidelines for biological properties that these functions should obey.

Yashin et al. (1985) are also focused on statistical inference rather than relatively qualitative descriptions and clarifications of logic, and as a result, make some rather restrictive assumptions about functional forms. For instance, in our language, Yashin et al., 1985 assume that the mortality rule *µ* must be quadratic in the failures (their section IV.E.2), whereas we make no such assumption, and indeed argue that a more biologically realistic mortality rule is not a finite degree polynomial (our ‘geometric’ mortality rule Eq. 13). Our Feynman-Kac representations also describe how precisely the stochastic mortality *µ* affects the dynamics of failure accumulation. In supplementary section S3, we are also able to obtain a direct correspondence (Eq. S30) between the probability densities of the unconditioned and the conditioned processes (our Eq. S7 and Eq. 4; Yashin et al.’s (1985) Eq. 5 and Eq. 7). By extending Woodbury and Manton’s (1977) and Yashin et al.’s (1985) model to more general mortality functions and connecting it to the damage/failure accumulation literature, we hope to bring attention to it and render it usable by a wider audience. The Feynman-Kac formulae also allow direct and efficient numerical predictions of expected mortality or failure curves under a given level of mortality/disappearance in terms of the much simpler Tithonus model Eq. 1.

Nielsen et al. (2024) have proposed a model they call the ‘multiple and inter-dependent component cause model’, where individuals are comprised of a large number of interdependent sub-systems, each capable of failure, arranged as a complete graph, and shown that their model predicts approximately Gompertz-Makeham mortality curves. Nielsen et al.’s (2024) model is the same as our logistic failure accumulation rule when our baseline failure rate is zero (*ϕ*_*ρ*_ = 0, compare their Eq. 3 with our Eq. 12; also see our supplementary section S6 for a derivation for a more general class of graphs). However, Nielsen et al. (2024) assume a linear mortality rule *µ*( *f* ) = *µ*_*e*_ + *χ f* and thus find late-life plateaus in mortality, whereas we advocate for a more realistic mortality rule (the ‘geometric mortality rule’ Eq. 13) on biological grounds. Nielsen et al. (2024) also work with our ‘Tithonus model’ rather than accounting for stochastic mortality and selective disappearance (to see this, note that their Eq. 9 is the Master equation obtained by setting *µ* ≡ 0 in Eq. S4 in supplementary section S1). In other words, their equations are not quite accurate unless deviations due to selective disappearance, as quantified by Eq. 8, are small enough to neglect.

On the evolutionary side, Bega and Hadany (2026) have recently studied how failure accumulation models interact with Hamilton’s (1966) selection shadow and its evolutionary consequences. Bega and Hadany’s (2026) starting point is an equation that is an alternative form of the ending point of our deterministic logistic failure accumulation with the geometric mortality rule. This can be made explicit by introducing a new variable *D*(*t*) := *f* (*t*)/(1 − *f* (*t*)) and re-expressing our Eq. 12 and Eq. 13 in terms of this new variable; doing so and setting *ϕ*_*ρ*_ = 0 yields the first set of ODEs in the leftmost box of Figure 1 of Bega and Hadany (2026) up to a change in notation. Importantly, Bega and Hadany (2026) assume the parameters *A* and *b* of a Gompertz curve of the form *µ*(*t*) = *Ae*^*bt*^ can evolve independently. For the Gompertz-Makeham curve that we have systematically derived in terms of more fundamental failure accumulation related parameters (Eq. 14), this *only* happens when either *ϕ*_*ρ*_ = 0 or *k*_*ρ*_ = 0, *i.e*. when either the ‘baseline failure rate’ of every sub-system is exactly zero or sub-systems have no effect on each other’s functioning. Since intra-organismal sub-systems are typically interdependent to at least some degree (*k*_*ρ*_ > 0) and have some risk of spontaneous failure due to stochastic wear-and-tear (*ϕ*_*ρ*_ > 0), we generally expect the Gompertz parameters in Eq. 14 to be correlated. Strehler and Mildvan (1960) and Gavrilov and Gavrilova (1991, sections 6.4-6.7) have also presented more mechanistically explicit reliability-theoretic models in which the Gompertz parameters are correlated when derived systematically from more fundamental underlying processes. At least for mortality curves arising from mechanistic failure accumulation models such as in Gavrilov and Gavrilova (1991) and our work, Bega and Hadany’s (2026) central assumption is hence a stringent one, and care is required in assessing when it is a reasonable assumption for any particular mechanistic model.

Weitz and Fraser (2001) have also put forth the idea that stochasticity followed by selective disappearance/mortality can cause late life plateaus without fixed differences in inter-individual quality. Though our models are somewhat similar, they differ in two crucial assumptions. Weitz and Fraser (2001) assume that failure accumulation (‘loss of vitality’ in their language) occurs at a constant rate (their Eq. 5; their constant *ε* plays the role of our function *r*( *f* ), compare with our Eq. 1). In contrast, in our work, we have argued that we should expect failure to beget failure, and the overall failure accumulation rate is thus non-constant (it is increasing). Weitz and Fraser (2001) also assume that an organism dies exactly when all its sub-systems have failed and no sooner (in their language, exactly when vitality hits 0), whereas we work with the more realistic assumption that organisms experience a continuous failure-dependent mortality risk.

### S9 Figure 4 for different values of *k*_*ρ*_ and *χ*

**Figure S2:**
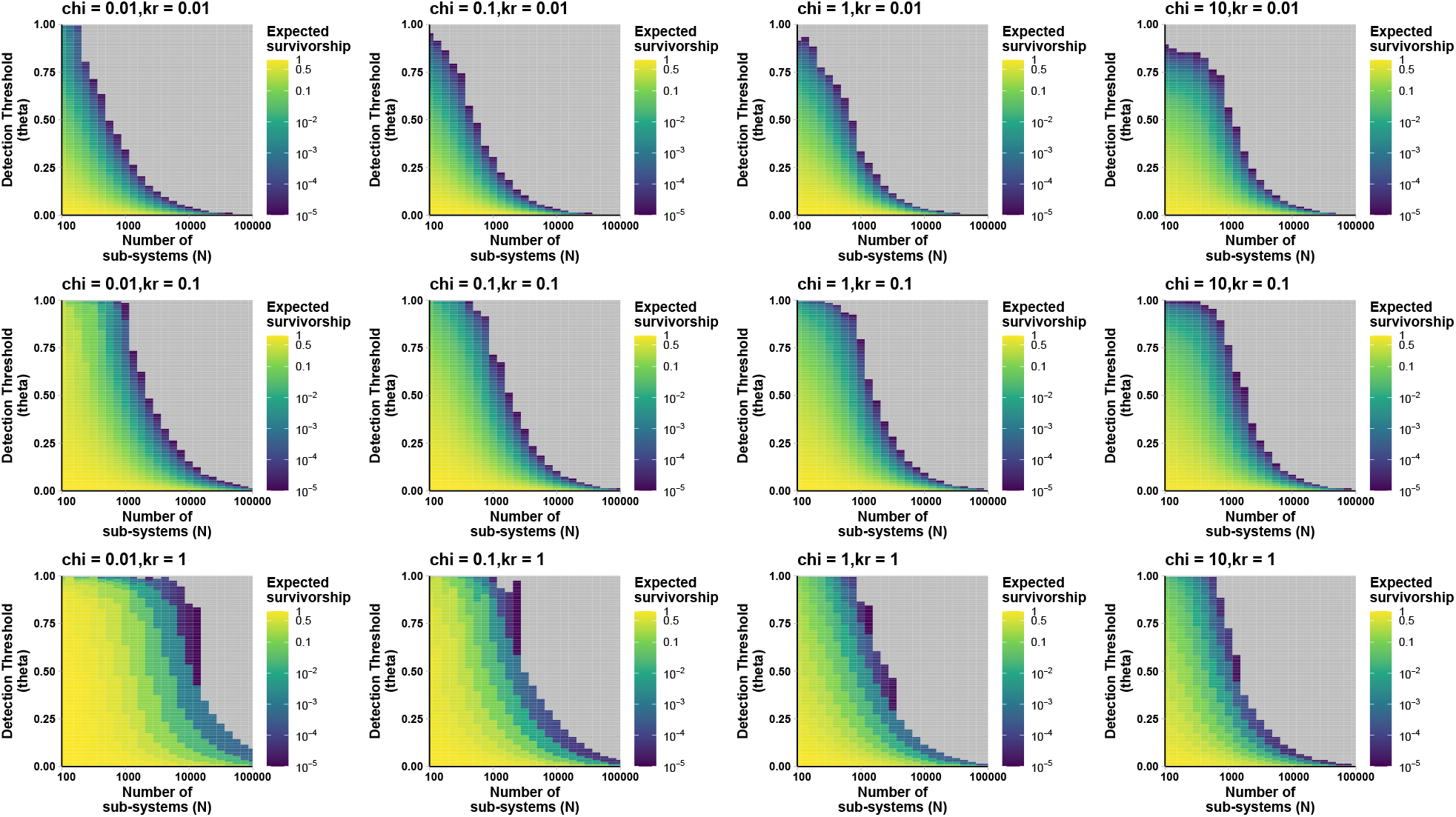
Effect of varying the interdependence (*k*_*ρ*_) and the impact of failures on mortality (*χ*) on the results presented in Fig 4. *k*_*ρ*_ increases along rows and *χ* increases across columns. Extrinsic mortality is *µ*_*e*_ = 10^−5^, and all other functional forms and parameter values are as in Fig 4.

Note that interdependence *k*_*ρ*_ controls how strongly failure affects failure rate, and *χ* controls how strongly failure affects mortality rate. Deviations from the Tithonus model are easiest to detect when failures impact failure rate (high *k*_*ρ*_) but not mortality rate (low *χ*) because selective disappearance can operate without greatly reducing the overall survivorship.

## Footnotes

1 A more rigorous statement of this approximation appears in Ethier and Kurtz, 1986, Chapter 11. The description is mathematically involved but quite standard (the technical phrase is ‘convergence of infinitesimal generators’), and we thus neglect mathematical precision here in favour of accessibility.

2 see Ethier and Kurtz, 1986 chapter 4 or Øksendal, 1998 section 7.3 for precise definitions of ‘nice’. In this work, we will assume all required regularity conditions are satisfied whenever such generic functions appear.

3 *i.e*. it vanishes at the boundaries of Ω.

4 in words, *M* and its first two derivatives should be continuous in [0, 1].

5 The reader may have noticed that in the supplement, this is the definition rather than the result Eq. 7 in the main text; we will derive that it is equivalent to the definition as the denominator of Eq. 6. We switch the order of definition and result here in the supplementary to make the intuition for deriving the result more transparent.

6 This is a slight abuse of notation, since *S*(f_*t*_ ) depends on the entire path {f_*s*_ }_*s*∈[0,*t*]_, and not just the final value f_*t*_.

7 This is the same as saying *M* is 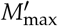-Lipschitz continuous. Using Feynman-Kac also requires *M* to be *C*^2^([0, 1]) (for Eq. S20).

8 Note that the same logic can be used to find an expression for a candidate *ρ*( *f* ) given a known mortality hazard *H*

9 because the fraction 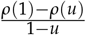 is a difference quotient that converges to *ρ*^′^(1) as *u* → 1

10 We have lost any potential signature of Makeham because we are working in the large *t* regime (and thus Gompertz dynamics dominate due to the exponential).

## Literature Cited

Allesina, S. (2026). “One Hundred Years of Lotka–Volterra Models”. Annual Review of Ecology, Evolution, and Systematics. doi: 10.1146/annurev-ecolsys-102924-040829.

Armitage, P. and Doll, R. (1954). “The age distribution of cancer and a multi-stage theory of carcinogenesis”. British Journal of Cancer 8.1, pp. 1–12. doi: 10.1038/sj.bjc.6602297.

Avila, P. and Lehmann, L. (2023). “Life history and deleterious mutation rate coevolution”. Journal of Theoretical Biology 573, p. 111598. doi: 10.1016/j.jtbi.2023.111598.

Avila, P., Priklopil, T., and Lehmann, L. (2021). “Hamilton’s rule, gradual evolution, and the optimal (feedback) control of phenotypically plastic traits”. Journal of Theoretical Biology 526, p. 110602. doi: 10.1016/j.jtbi.2021.110602.

Balan, T. A. and Putter, H. (2020). “A tutorial on frailty models”. Statistical Methods in Medical Research 29.11, pp. 3424–3454. doi: 10.1177/0962280220921889.

Barbi, E., Lagona, F., Marsili, M., Vaupel, J. W., and Wachter, K. W. (2018). “The plateau of human mortality: Demography of longevity pioneers”. Science 360.6396, pp. 1459–1461. doi: 10.1126/science.aat3119.

Baudisch, A. (2012). “Birds Do It, Bees Do It, We Do It: Contributions of Theoretical Modelling to Understanding the Shape of Ageing across the Tree of Life”. Gerontology 58.6, pp. 481–489. doi: 10.1159/000341861.

Bega, D. and Hadany, L. (2026). “Evolution of Senescence by Damage Accumulation That Accelerates With Age Throughout an Organism’s Lifespan”. Ecology and Evolution 16.3. doi: 10.1002/ece3.72988.

Belikov, A. V. (2019). “Age-related diseases as vicious cycles”. Ageing Research Reviews 49, pp. 11–26. doi: 10.1016/j.arr.2018.11.002.

Bernard, C., Compagnoni, A., and Salguero-Gómez, R. (2020). “Testing Finch’s hypothesis: The role of organismal modularity on the escape from actuarial senescence”. Functional Ecology 34.1, pp. 88–106. doi: 10.1111/1365-2435.13486.

Bhat, A. S. (2025). “A stochastic field theory for the evolution of quantitative traits in finite populations”. Theoretical Population Biology 161, pp. 1–12. doi: 10.1016/j.tpb.2024.10.003.

Bhat, A. S. and Guttal, V. (2025). “Eco-Evolutionary Dynamics for Finite Populations and the Noise-Induced Reversal of Selection”. The American Naturalist 205.1, pp. 1–19. doi: 10.1086/733196.

Bhat, A. S. and Kokko, H. (submitted). “Demographic senescence as multi-level selection in miniature”. In Review, Population Biology Modeling & Theory. Available on aRxiv. doi: 10.48550/arXiv.2607.00262.

Boonekamp, J. J., Briga, M., and Verhulst, S. (2015). “The heuristic value of redundancy models of aging”. Experimental Gerontology 71, pp. 95–102. doi: 10.1016/j.exger.2015.09.005.

Bouwhuis, S., Sheldon, B., Verhulst, S., and Charmantier, A. (2009). “Great tits growing old: selective disappearance and the partitioning of senescence to stages within the breeding cycle”. Proceedings of the Royal Society B: Biological Sciences 276.1668, pp. 2769–2777. doi: 10.1098/rspb.2009.0457.

Camarda, C. G. (2022). “The curse of the plateau. Measuring confidence in human mortality estimates at extreme ages”. Theoretical Population Biology 144, pp. 24–36. doi: 10.1016/j.tpb.2022.01.002.

Carey, J. R., Liedo, P., Orozco, D., and Vaupel, J. W. (1992). “Slowing of Mortality Rates at Older Ages in Large Medfly Cohorts”. Science 258.5081, pp. 457–461. doi: 10.1126/science.1411540.

Chmilar, S. L., Luzardo, A. C., Dutt, P., Pawluk, A., Thwaites, V. C., and Laird, R. A. (2024). “Caloric restriction extends lifespan in a clonal plant”. Ecology Letters 27.6, e14444. doi: 10.1111/ele.14444.

Cohen, A. A., Coste, C. F. D., Li, X.-Y., Bourg, S., and Pavard, S. (2020). “Are trade-offs really the key drivers of ageing and life span?” Functional Ecology 34.1, pp. 153–166. doi: 10.1111/1365-2435.13444.

Cohen, A. A., Ferrucci, L., Fülöp, T., Gravel, D., Hao, N., Kriete, A., Levine, M. E., Lipsitz, L. A., Olde Rikkert, M. G. M., Rutenberg, A., Stroustrup, N., and Varadhan, R. (2022). “A complex systems approach to aging biology”. Nature Aging 2.7, pp. 580–591. doi: 10.1038/s43587-022-00252-6.

Cooney, D. B., Levin, S. A., Mori, Y., and Plotkin, J. B. (2023). “Evolutionary dynamics within and among competing groups”. Proceedings of the National Academy of Sciences 120.20, e2216186120. doi: 10.1073/pnas.2216186120.

Curtsinger, J. W., Fukui, H. H., Townsend, D. R., and Vaupel, J. W. (1992). “Demography of Genotypes: Failure of the Limited Life-Span Paradigm in Drosophila melanogaster”. Science 258.5081, pp. 461–463. doi: 10.1126/science.1411541.

Farrell, S., Stubbings, G., Rockwood, K., Mitnitski, A., and Rutenberg, A. (2021). “The potential for complex computational models of aging”. Mechanisms of Ageing and Development 193, p. 111403. doi: 10.1016/j.mad.2020.111403.

Feehan, D. M. (2018). “Separating the Signal From the Noise: Evidence for Deceleration in Old-Age Death Rates”. Demography 55.6, pp. 2025–2044. doi: 10.1007/s13524-018-0728-x.

Feynman, R. P. (1948). “Space-Time Approach to Non-Relativistic Quantum Mechanics”. Rev. Mod. Phys. 20 (2), pp. 367–387. doi: 10.1103/RevModPhys.20.367.

Finch, C. E. (1994). Longevity, Senescence, and the Genome. The John D. and Catherine T. MacArthur Foundation Series on Mental Health and Development. Chicago, IL: University of Chicago Press. ISBN: 978-0-226-24889-9.

Finch, C. E. (2009). “Update on Slow Aging and Negligible Senescence – A Mini-Review”. Gerontology 55.3, pp. 307–313. doi: 10.1159/000215589.

Finkelstein, M. and Esaulova, V. (2006). “Asymptotic behavior of a general class of mixture failure rates”. Advances in Applied Probability 38.1, pp. 244–262. doi: 10.1239/aap/1143936149.

Fontanari, J. F. and Serva, M. (2014). “Nonlinear group survival in Kimura’s model for the evolution of altruism”. Mathematical Biosciences 249, pp. 18–26. doi: 10.1016/j.mbs.2014.01.003.

Fox, C. W. and Moya-Laraño, J. (2003). “Why organisms show late-life mortality plateaus: a null model for comparing patterns of mortality”. Evolutionary Ecology Research 5.7, pp. 999–1009.

Gavrilov, L. A. and Gavrilova, N. S. (1991). The Biology of Life Span: A Quantitative Approach. Harwood Academic Publishers. ISBN: 978-3-7186-4983-9.

Gavrilov, L. A. and Gavrilova, N. S. (2001). “The Reliability Theory of Aging and Longevity”. Journal of Theoretical Biology 213.4, pp. 527–545. doi: 10.1006/jtbi.2001.2430.

Gavrilov, L. A. and Gavrilova, N. S. (2006). “Models of Systems Failure in Aging”. In: Handbook of Models for Human Aging. Ed. by P. Michael Conn. Elsevier, pp. 45–67. ISBN: 978-0-12-369391-4. doi: 10.1016/b978-012369391-4/50006-0.

Gavrilov, L. A. and Gavrilova, N. S. (2019). “Late-life mortality is underestimated because of data errors”. PLOS Biology 17.2, e3000148. doi: 10.1371/journal.pbio.3000148.

Gillespie, D. T. (1977). “Exact stochastic simulation of coupled chemical reactions”. The Journal of Physical Chemistry 81.25, pp. 2340–2361. doi: 10.1021/j100540a008.

Gorbunova, V., Seluanov, A., Mao, Z., and Hine, C. (2007). “Changes in DNA repair during aging”. Nucleic Acids Research 35.22, pp. 7466–7474. doi: 10.1093/nar/gkm756.

Graves, R. (2017). The Greek Myths: The Complete and Definitive Edition. Penguin Books Limited. ISBN: 9780241983386.

Hämäläinen, A., Dammhahn, M., Aujard, F., Eberle, M., Hardy, I., Kappeler, P. M., Perret, M., Schliehe-Diecks, S., and Kraus, C. (2014). “Senescence or selective disappearance? Age trajectories of body mass in wild and captive populations of a small-bodied primate”. Proceedings of the Royal Society B: Biological Sciences 281.1791, p. 20140830. doi: 10.1098/rspb.2014.0830.

Hamilton, W. D. (1966). “The moulding of senescence by natural selection”. Journal of Theoretical Biology 12.1, pp. 12–45. doi: 10.1016/0022-5193(66)90184-6.

Hayward, A. D., Wilson, A. J., Pilkington, J. G., Clutton-Brock, T. H., Pemberton, J. M., and Kruuk, L. E. B. (2013). “Reproductive senescence in female Soay sheep: variation across traits and contributions of individual ageing and selective disappearance”. Functional Ecology 27.1, pp. 184–195. doi: 10.1111/1365-2435.12029.

Itô, K. (1951). “On a formula concerning stochastic differentials”. Nagoya Mathematical Journal 3, pp. 55–65. doi: 10.1017/S0027763000012216.

Kac, M. (1949). “On distributions of certain Wiener functionals”. Transactions of the American Mathematical Society 65.1, pp. 1–13. doi: 10.2307/1990512.

Karlin, S. and Tavaré, S. (1982). “A diffusion process with killing: the time to formation of recurrent deleterious mutant genes”. Stochastic Processes and their Applications 13.3, pp. 249–261. doi: 10.1016/0304-4149(82)90012-6.

Karlin, S. and Taylor, H. M. (1981). A second course in stochastic processes. New York: Academic Press. ISBN: 978-0-12-398650-4.

Kimura, M. (1984). “Evolution of an Altruistic Trait through Group Selection as Studied by the Diffusion Equation Method”. IMA Journal of Mathematics Applied in Medicine & Biology 1.1, pp. 1–15. doi: 10.1093/imammb/1.1.1.

Kirkwood, T. B. L. (1977). “Evolution of ageing”. Nature 270.5635, pp. 301–304. doi: 10.1038/270301a0.

Kokko, H. (2024). “Who is afraid of modelling time as a continuous variable?” Methods in Ecology and Evolution 15.10, pp. 1736–1756. doi: 10.1111/2041-210X.14394.

Komarova, N. L. and Wodarz, D. (2025). “Population structure reverses selection of variants with proportionally scaled birth and death rates”. Nature Communications 17.1. doi: 10.1038/s41467-025-66951-x.

Laird, R. A. and Sherratt, T. N. (2009). “The evolution of senescence through decelerating selection for system reliability”. Journal of Evolutionary Biology 22.5, pp. 974–982. doi: 10.1111/j.1420-9101.2009.01709.x.

Laird, R. A. and Sherratt, T. N. (2010). “The evolution of senescence in multi-component systems”. Biosystems 99.2, pp. 130–139. doi: 10.1016/j.biosystems.2009.10.008.

Ledberg, A. (2020). “Exponential increase in mortality with age is a generic property of a simple model system of damage accumulation and death”. PLOS ONE 15.6, e0233384. doi: 10.1371/journal.pone.0233384.

Lehtonen, J. (2020). “Longevity and the drift barrier: Bridging the gap between Medawar and Hamilton”. Evolution Letters 4.4, pp. 382–393. doi: 10.1002/evl3.173.

López-Otín, C., Blasco, M. A., Partridge, L., Serrano, M., and Kroemer, G. (2023). “Hallmarks of aging: An expanding universe”. Cell 186.2, pp. 243–278. doi: 10.1016/j.cell.2022.11.001.

Luo, S. (2014). “A unifying framework reveals key properties of multilevel selection”. Journal of Theoretical Biology 341, pp. 41–52. doi: 10.1016/j.jtbi.2013.09.024.

Maklakov, A. A. and Chapman, T. (2019). “Evolution of ageing as a tangle of trade-offs: energy versus function”. Proceedings of the Royal Society B: Biological Sciences 286.1911, p. 20191604. doi: 10.1098/rspb.2019.1604.

Medawar, P. B. (1952). An Unsolved Problem of Biology: An Inaugural Lecture Delivered at University College, London, 6 December, 1951. H.K. Lewis and Company.

Meyer, D. H., Maklakov, A. A., and Schumacher, B. (2025). “Aging by the clock and yet without a program”. Nature Aging, pp. 1–11. doi: 10.1038/s43587-025-00975-2.

Meyer, D. H. and Schumacher, B. (2024). “Aging clocks based on accumulating stochastic variation”. Nature Aging 4.6, pp. 871–885. doi: 10.1038/s43587-024-00619-x.

Montanari, A. N., Zanin, P., and Motter, A. E. (2026). “Disorder-promoted stability”. Science 393.6817, pp. 1241–1249. doi: 10.1126/science.aeg3946.

Newman, S. J. (2018). “Errors as a primary cause of late-life mortality deceleration and plateaus”. PLOS Biology 16.12, e2006776. doi: 10.1371/journal.pbio.2006776.

Nielsen, P. Y., Jensen, M. K., Mitarai, N., and Bhatt, S. (2024). “The Gompertz Law emerges naturally from the inter-dependencies between sub-components in complex organisms”. Scientific Reports 14.1, p. 1196. doi: 10.1038/s41598-024-51669-5.

Øksendal, B. K. (1998). Stochastic differential equations: an introduction with applications. Springer. ISBN: 978-3-662-03620-4.

Pichler, A. and Uhlig, D. (2023). “Mortality in Germany during the COVID-19 Pandemic”. International Journal of Environmental Research and Public Health 20.20, p. 6942. doi: 10.3390/ijerph20206942.

Pletcher, S. D. and Curtsinger, J. W. (1998). “Mortality Plateaus and the Evolution of Senescence: Why are Oldage Mortality Rates so Low?” Evolution 52.2, pp. 454–464. doi: 10.1111/j.1558-5646.1998.tb01645.x.

Rosen, R. (1958). “A relational theory of biological systems”. The Bulletin of Mathematical Biophysics 20.3, pp. 245–260. doi: 10.1007/BF02478302.

Shefferson, R. P., Jones, O. R., and Salguero-Gómez, R. (2017). The Evolution of Senescence in the Tree of Life. Cambridge University Press. ISBN: 978-1-107-07850-5.

Siler, W. (1979). “A Competing-Risk Model for Animal Mortality”. Ecology 60.4, pp. 750–757. doi: 10.2307/1936612.

Snyder, R. E., Ellner, S. P., and Hooker, G. (2021). “Time and Chance: Using Age Partitioning to Understand How Luck Drives Variation in Reproductive Success”. The American Naturalist 197.4, E110–E128. doi: 10.1086/712874.

Steinsaltz, D. and Evans, S. (2004). “Markov mortality models: implications of quasistationarity and varying initial distributions”. Theoretical Population Biology 65.4, pp. 319–337. doi: 10.1016/j.tpb.2003.10.007.

Steinsaltz, D. and Evans, S. (2006). “Quasistationary distributions for one-dimensional diffusions with killing”. Transactions of the American Mathematical Society 359.3, pp. 1285–1324. doi: 10.1090/s0002-9947-06-03980-8.

Strehler, B. L. and Mildvan, A. S. (1960). “General Theory of Mortality and Aging”. Science 132.3418, pp. 14–21. doi: 10.1126/science.132.3418.14.

Sun, E. D., Michaels, T. C. T., and Mahadevan, L. (2020). “Optimal control of aging in complex networks”. Proceedings of the National Academy of Sciences 117.34, pp. 20404–20410. doi: 10.1073/pnas.2006375117.

Tian, Y. E., Cropley, V., Maier, A. B., Lautenschlager, N. T., Breakspear, M., and Zalesky, A. (2023). “Hetero-geneous aging across multiple organ systems and prediction of chronic disease and mortality”. Nature Medicine 29.5, pp. 1221–1231. doi: 10.1038/s41591-023-02296-6.

Tuljapurkar, S., Steiner, U. K., and Orzack, S. H. (2009). “Dynamic heterogeneity in life histories”. Ecology Letters 12.1, pp. 93–106. doi: 10.1111/j.1461-0248.2008.01262.x.

Vaupel, J. W., Baudisch, A., Dölling, M., A. Roach D., and Gampe, J. (2004). “The case for negative senescence”. Theoretical Population Biology. Demography in the 21st Century 65.4, pp. 339–351. doi: 10.1016/j.tpb.2003.12.003.

Vaupel, J. W., Carey, J. R., Christensen, K., Johnson, T. E., Yashin, A. I., Holm, N. V., Iachine, I. A., Kannisto, V., Khazaeli, A. A., Liedo, P., Longo, V. D., Zeng, Y., Manton, K. G., and Curtsinger, J. W. (1998). “Biodemographic Trajectories of Longevity”. Science 280.5365, pp. 855–860. doi: 10.1126/science.280.5365.855.

Vaupel, J. W., Manton, K. G., and Stallard, E. (1979). “The impact of heterogeneity in individual frailty on the dynamics of mortality”. Demography 16.3, pp. 439–454. doi: 10.2307/2061224.

Vaupel, J. W. and Missov, T. (2014). “Unobserved population heterogeneity: A review of formal relationships”. Demographic Research 31, pp. 659–686. doi: 10.4054/DemRes.2014.31.22.

De Vries, C., Galipaud, M., and Kokko, H. (2023). “Extrinsic mortality and senescence: a guide for the perplexed”. Peer Community Journal 3. doi: 10.24072/pcjournal.253.

Vural, D. C., Morrison, G., and Mahadevan, L. (2014). “Aging in complex interdependency networks”. Physical Review E 89.2, p. 022811. doi: 10.1103/PhysRevE.89.022811.

Webster, A. J. (2019). “Multi-stage models for the failure of complex systems, cascading disasters, and the onset of disease”. PLOS ONE 14.5, e0216422. doi: 10.1371/journal.pone.0216422.

Weitz, J. S. and Fraser, H. B. (2001). “Explaining mortality rate plateaus”. Proceedings of the National Academy of Sciences 98.26, pp. 15383–15386. doi: 10.1073/pnas.261228098.

Williams, G. C. (1957). “Pleiotropy, Natural Selection, and the Evolution of Senescence”. Evolution 11.4, pp. 398–411. doi: 10.1111/j.1558-5646.1957.tb02911.x.

Williams, G. C. (1999). “The Tithonus Error in Modern Gerontology”. The Quarterly Review of Biology 74.4, pp. 405–415. doi: 10.1086/394111.

Woodbury, M. A. and Manton, K. G. (1977). “A random-walk model of human mortality and aging”. Theoretical Population Biology 11.1, pp. 37–48. doi: 10.1016/0040-5809(77)90005-3.

Woodbury, M. A. and Manton, K. G. (1983). “A Mathematical Model of the Physiological Dynamics of Aging and Correlated Mortality Selection. I. Theoretical Development and Critiques”. Journal of Gerontology 38.4, pp. 398–405. doi: 10.1093/geronj/38.4.398.

Yashin, A. I., Manton, K. G., and Vaupel, J. W. (1985). “Mortality and aging in a heterogeneous population: A stochastic process model with observed and unobserved variables”. Theoretical Population Biology 27.2, pp. 154–175. doi: 10.1016/0040-5809(85)90008-5.

Yashin, A. I., Vaupel, J. W., and Iachine, I. A. (1994). “A duality in aging: the equivalence of mortality models based on radically different concepts”. Mechanisms of Ageing and Development 74.1, pp. 1–14. doi: 10.1016/0047-6374(94)90094-9.

## References Cited Only in the Supplementary

Black, A. J. and McKane, A. J. (2012). “Stochastic formulation of ecological models and their applications”. Trends in Ecology & Evolution 27.6, pp. 337–345. doi: 10.1016/j.tree.2012.01.014.

Ethier, S. N. and Kurtz, T. G. (1986). Markov Processes: Characterization and Convergence. Wiley. ISBN: 9780470316658. doi: 10.1002/9780470316658.

Godsil, C. and Royle, G. (2001). Algebraic graph theory. Graduate texts in mathematics. Springer. ISBN: 978-0-387-95220-8.

Karatzas, I. and Shreve, S. E. (1998). Brownian motion and stochastic calculus. Graduate texts in mathematics. Springer. ISBN: 978-0-387-97655-6.

McAvoy, A. and Hauert, C. (2015). “Structural symmetry in evolutionary games”. Journal of The Royal Society Interface 12.111, p. 20150420. doi: 10.1098/rsif.2015.0420.

## Literature Cited in the Supplement

Gardiner, C. W. (2009). Stochastic methods: a handbook for the natural and social sciences. Berlin: Springer. ISBN: 978-3-540-70712-7.

Gavrilov, L. A. and Gavrilova, N. S. (1991). The Biology of Life Span: A Quantitative Approach. Harwood Academic Publishers. isbn: 978–3-7186-4983-9.

Manton, K. G., Woodbury, M. A., and Stallard, E. (1988). “Models of the interaction of mortality and the evolution of risk factor distribution: A general stochastic process formulation”. Statistics in Medicine 7. 1-2, pp. 239–256. doi: 10.1002/sim.4780070125.

Mulder, P. G. H. (1993). “The simultaneous processes of ageing and mortality”. Statistica Neerlandica 47.4, pp. 253–267. doi: 10.1111/j.1467-9574.1993.tb01422.x.

Woodbury, M. A. and Manton, K. G.(1983). “A Mathematical Model of the Physiological Dynamics of Aging and Correlated Mortality Selection. I. Theoretical Development and Critiques”. Journal of Gerontology 38.4, pp. 398–405. doi: 10.1093/geronj/38.4.398.

